# Deubiquitinase-Targeting Chimeras for Targeted Protein Stabilization

**DOI:** 10.1101/2021.04.30.441959

**Authors:** Nathaniel J. Henning, Lydia Boike, Jessica N. Spradlin, Carl C. Ward, Bridget Belcher, Scott M. Brittain, Matthew Hesse, Dustin Dovala, Lynn M. McGregor, Jeffrey M. McKenna, John A. Tallarico, Markus Schirle, Daniel K. Nomura

## Abstract

Targeted protein degradation is a powerful therapeutic modality that uses heterobifunctional small-molecules to induce proximity between E3 ubiquitin ligases and target proteins to ubiquitinate and degrade specific proteins of interest. However, many proteins are ubiquitinated and degraded to drive disease pathology; in these cases targeted protein stabilization (TPS), rather than degradation, of the actively degraded target using a small-molecule would be therapeutically beneficial. Here, we present the Deubiquitinase-Targeting Chimera (DUBTAC) platform for TPS of specific proteins. Using chemoproteomic approaches, we discovered the covalent ligand EN523 that targets a non-catalytic allosteric cysteine C23 in the K48 ubiquitin-specific deubiquitinase OTUB1. We then developed a heterobifunctional DUBTAC consisting of our EN523 OTUB1 recruiter linked to lumacaftor, a drug used to treat cystic fibrosis that binds ΔF508-CFTR. We demonstrated proof-of-concept of TPS by showing that this DUBTAC robustly stabilized ΔF508-CFTR in human cystic fibrosis bronchial epithelial cells in an OTUB1-dependent manner. Our study underscores the utility of chemoproteomics-enabled covalent ligand discovery approaches to develop new induced proximity-based therapeutic modalities and introduces the DUBTAC platform for TPS.

**Editorial summary:** We have developed the Deubiquitinase Targeting Chimera (DUBTAC) platform for targeted protein stabilization. We have discovered a covalent recruiter against the deubiquitinase OTUB1 that we have linked to the mutant ΔF508-CFTR targeting cystic fibrosis drug Lumacaftor to stabilize mutant CFTR protein in cells.

## Main

One of the biggest challenges in developing new disease therapies is that most, >85 %, of the proteome is considered “undruggable,” meaning the majority of proteins are devoid of well-characterized, functional binding pockets or “ligandable hotspots” that small-molecules can bind to modulate the protein’s function ^1,2^. Engaging the mostly undruggable proteome to uncover new disease therapies not only requires technological innovations that facilitate rapid discovery of ligandable hotspots across the proteome, but also demands new therapeutic modalities that alter protein function through novel mechanisms. Targeted protein degradation (TPD) has arisen as a powerful therapeutic modality for tackling the undruggable proteome by targeting specific proteins for ubiquitination and proteasomal degradation. The small-molecule effectors of TPD, called proteolysis-targeting chimeras (PROTACs), employ heterobifunctional molecules that consist of an E3 ligase recruiter linked to a protein-targeting ligand to induce the formation of ternary complexes that bring together an E3 ubiquitin ligase and the target protein as a neo-substrate ^3–5^. This results in the polyubiquitination and degradation of the target, which enables targeting otherwise intractable proteins without functional binding sites. New approaches for TPD have also arisen that exploit endosomal and lysosomal degradation pathways with Lysosome Targeting Chimeras (LYTACs) or autophagy with Autophagy Targeting Chimeras (AUTACs) ^6,7^. While TPD with PROTACs, LYTACs, or AUTACs enables the targeted degradation of potentially any protein target within, at the surface, or outside of the cell, these induced-proximity approaches are limited to degradation of proteins. New approaches for chemically induced proximity beyond degradation have also been developed in recent years, including targeted phosphorylation with Phosphorylation-Inducing Chimeric Small-Molecules (PHICS) and targeted dephosphorylation, but no small-molecule based induced proximity approaches exist for targeted deubiquitination ^8,9^.

The active ubiquitination and degradation of proteins is the root cause of several classes of diseases, including many tumor suppressors in cancer (e.g. TP53, CDKN1A, CDN1C, BAX), mutated and misfolded proteins such as ΔF508-CFTR in cystic fibrosis, or glucokinase in pancreatic cells, which would benefit from a targeted protein stabilization (TPS) therapeutic strategy, rather than degradation ^10–13^. Analogous to TPD, we hypothesized that TPS would be enabled by the discovery of a small-molecule recruiter of a deubiquitinase (DUB) that could be linked to a protein-targeting ligand to form a chimeric molecule which would induce the deubiquitination and stabilization of proteins of interest—a heterobifunctional stabilizer or Deubiquitinase Targeting Chimera (DUBTAC) **(Figure 1a)**. In this study, we report the discovery of a covalent ligand for the K48 ubiquitin-chain specific DUB OTUB1 which when linked to a protein-targeting ligand is able to stabilize a target protein to demonstrate proof-of-concept for the DUBTAC platform.

**Figure 1.**
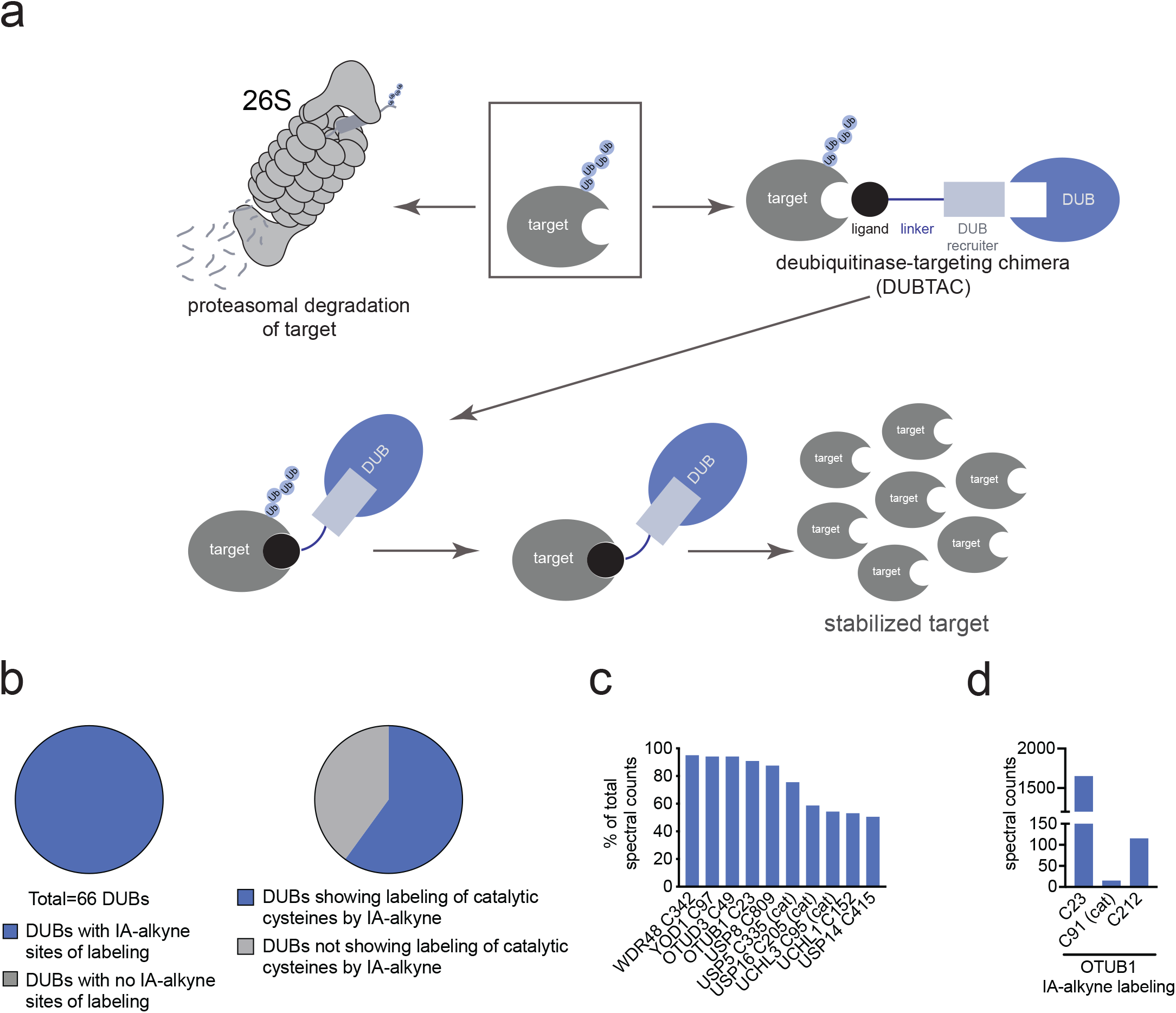
DUBTAC platform. **(a)** DUBTACs consist of heterobifunctional molecules linking a protein-targeting ligand to a DUB recruiter to induce proximity between a DUB and a ubiquitinated target protein to deubiquitinate and stabilize the levels of the protein. **(b)** Out of 65 DUBs mined in our research group’s aggregate chemoproteomic datasets of cysteine-reactive probe labeling with IA-alkyne in various complex proteomes, we identified probe-modified cysteines across all 100 % of the 65 DUBs. This is shown in the first pie chart. Among the 65 DUBs that showed probe-modified cysteines, 39 of these DUBs showed >10 aggregate spectral counts across our chemoproteomic datasets. 24 DUBs, or 62 %, of these 39 DUBs showed labeling of the DUB catalytic or active site cysteines. **(c)** Mining the DUB data, we identified 10 DUBs wherein there was one probe-modified cysteine that represented >50 % of the total aggregate spectral counts for probe-modified cysteine peptides for the particular DUB. 7 of those 10 DUBs do not target a known catalytic cysteine whereas 3 do target the catalytic cysteine (abbreviated by cat). **(d)** Analysis of aggregate chemoproteomic data for OTUB1 IA-alkyne labeling showing that C23 is the dominant site labeled by IA-alkyne compared to the catalytic (cat) C91. Chemoproteomic data analysis of DUBs across aggregated datasets can be found in **Table S1.**

To enable the DUBTAC platform, our first goal was to identify a small-molecule DUB recruiter that targets an allosteric site and did not inhibit DUB function, since the recruitment of a functional DUB would be required to deubiquitinate and stabilize the levels of a target protein. While many DUBs possess well-defined active sites bearing a catalytic and highly nucleophilic cysteine, there have not yet been systematic evaluations of allosteric, non-catalytic, and ligandable sites on DUBs that could be pharmacologically targeted to develop a DUB recruiter. Chemoproteomic platforms such as activity-based protein profiling (ABPP) have proven to be powerful approaches to map proteome-wide covalently ligandable sites directly in complex proteomes. ABPP utilizes reactivity-based amino acid-specific chemical probes to profile proteome-wide reactive, functional, and potentially ligandable sites directly in complex proteomes ^2,14^. When used in a competitive manner, covalent small-molecule ligands can be competed against probe binding to recombinant protein, in complex proteomes, in living cells, or in whole animals to enable covalent ligand discovery against potential ligandable sites revealed by reactivity-based probes ^2,15^. Previous studies have shown that isotopic tandem orthogonal proteolysis-ABPP (isoTOP-ABPP) platforms for mapping sites of labeling with reactivity-based probes using quantitative proteomic approaches can identify hyper-reactive, functional, and ligandable cysteines ^14,15^. To identify DUB candidates that possess potential ligandable allosteric cysteines, we analyzed our research group’s aggregate chemoproteomic data of proteome-wide sites modified by reactivity-based probes collected since the start of our laboratory. Specifically, we mined our collective chemoproteomic data of cysteine-reactive alkyne-functionalized iodoacetamide (IA-alkyne) probe labeling sites from isoTOP-ABPP experiments from 455 distinct experiments in human cell line proteomes for total aggregate spectral counts identified for each probe-modified site across the DUB family. We postulated that probe-modified cysteines within DUBs that showed the highest spectral counts aggregated over all chemoproteomic datasets, compared to those sites within the same DUB that showed lower spectral counts, may represent more reactive and potentially more ligandable cysteines. Caveats to this premise include cysteines that might be located in regions within a protein sequence that does not yield suitable tryptic peptides with respect to ionization and compatibility with mass spectrometry-based sequencing, and labeling of surface-exposed cysteines that may not be part of binding pockets.

However, we conjectured that the aggregate chemoproteomics data would still yield candidate allosteric ligandable sites within DUBs that could be prioritized for covalent ligand screening. We initially mined our aggregate chemoproteomic data for 66 members of the cysteine protease family of DUBs—including ubiquitin-specific proteases (USPs), ubiquitin C-terminal hydrolases (UCHs), Machado-Josephin domain proteases (MJDs) and ovarian tumour proteases (OTU)—since they encompass the majority of DUB superfamilies. Interestingly, we found probe-modified cysteines in 100 % of these DUB enzymes **(Figure 1b; Table S1).** Consistent with our aggregate chemoproteomic data of probe-modified sites being enriched in functional sites within DUBs, among the 40 DUBs that showed a total of >10 aggregate spectral counts of probe-modified peptides, 24 of those DUBs, 60 %, showed labeling of the DUB catalytic cysteine **(Figure 1b)**. We next prioritized this list of 40 DUBs to identify candidates wherein the dominant probe-modified cysteine was: 1) not the catalytic cysteine since we wanted to identify a non-catalytic allosteric site that we could target with a covalent ligand while retaining the catalytic activity; 2) in a dominantly identified probe-labeled peptide compared to other probe-modified sites within the same DUB, which - even without correcting for compatibility with MS/MS-based peptide identification - would indicate a high degree of reactivity and hopefully covalent ligandability of the particular allosteric cysteine compared to the catalytic site; and 3) frequently identified in chemoproteomics datasets as this would indicate the general accessibility of the cysteine in complex proteomes. We found 10 DUBs where one probe-modified cysteine represented >50 % of the spectral counts of all modified cysteines for the particular protein, of which 7 of these DUBs showed primary sites of probe-modification that did not correspond to the catalytic cysteine **(Figure 1c)**. Of these 10 DUBs, OTUB1 C23 was captured with >1000 total aggregate spectral counts compared to <500 aggregate spectral counts for the other DUBs **(Extended Data Figure 1a)**. In our aggregated chemoproteomic data, the tryptic peptide encompassing OTUB1 C23 was the dominant peptide labeled by IA-alkyne with >1500 total spectral counts, compared to 15 spectral counts for the peptide encompassing the catalytic C91, and 115 spectral counts for C212 **(Figure 1d)**. OTUB1 is a highly expressed DUB that specifically cleaves K48-linked polyubiquitin chains—the type of ubiquitin linkage that destines proteins for proteasome-mediated degradation—and C23 represents a known ubiquitin substrate recognition site that is distant from the catalytic C91 ^16–19^. Given our analysis of chemoproteomic data, the properties of OTUB1, and the location of C23, we chose OTUB1 as our candidate DUB for covalent ligand screening using gel-based ABPP approaches to discover an OTUB1 recruiter.

We performed a gel-based ABPP screen in which we screened 702 cysteine-reactive covalent ligands against labeling of pure OTUB1 protein with a rhodamine-functionalized cysteine-reactivity iodoacetamide (IA-rhodamine) probe **(Figure 2a; Extended Data Figure 1b; Table S2).** Through this screen, we identified the acrylamide EN523 as a top hit **(Figure 2b)**. We confirmed that EN523 dose-responsively displaced IA-rhodamine labeling of OTUB1 without causing any protein aggregation or precipitation **(Figure 2c).** We next performed liquid chromatography-tandem mass spectrometry analysis (LC-MS/MS) of tryptic peptides from EN523 bound to OTUB1 and showed that EN523 selectively targets C23, with no detectable modification of the catalytic C91 **(Figure 2d)**. Following these data, we performed an *in vitro* reconstituted OTUB1 deubiquitination activity assay monitoring monoubiquitin release from di-ubiquitin in the presence of OTUB1-stimulating UBE2D1, and demonstrated that EN523 does not inhibit OTUB1 deubiquitination activity **(Figure 2e)** ^20^.

**Figure 2.**
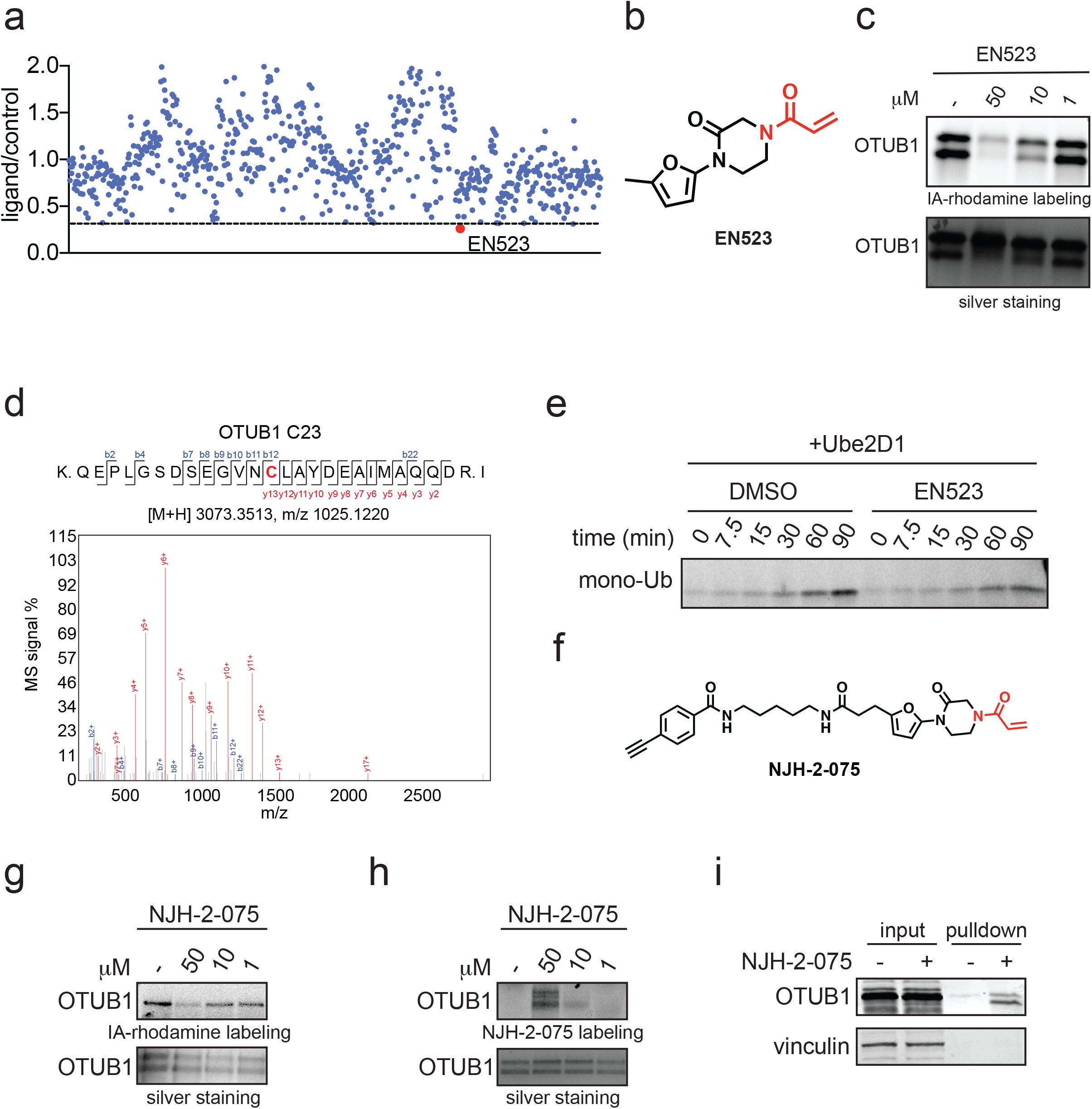
Discovery of covalent ligands that target OTUB1. **(a)** Covalent ligand screen of a cysteine-reactive library competed against IA-rhodamine labeling of recombinant OTUB1 to identify binders to OTUB1 by gel-based ABPP. Vehicle DMSO or cysteine-reactive covalent ligands (50 μM) were pre-incubated with OTUB1 for 30 min at room temperature prior to IA-rhodamine labeling (500 nM, 30 min room temperature). OTUB1 was then separated by SDS/PAGE and in-gel fluorescence was assessed and quantified. Gel-based ABPP data and quantification of in-gel fluorescence shown in **Extended Data Figure 1b** and **Table S2**. EN523 annotated in red was the top hit that showed the greatest inhibition of OTUB1 IA-rhodamine labeling. **(b)** Structure of EN523 shown with cysteine-reactive acrylamide highlighted in red. **(c)** Gel-based ABPP confirmation showing dose-responsive inhibition of IA-rhodamine binding of OTUB1. Vehicle DMSO or EN523 were pre-incubated with OTUB1 for 30 min at 37 °C prior to IA-rhodamine labeling (500 nM, 30 min room temperature). OTUB1 was then separated by SDS/PAGE and in-gel fluorescence was assessed. Also shown is silver staining showing protein loading. Shown is a representative gel of n=3 biologically independent samples/group. **(d)** LC-MS/MS data showing EN523-modified adduct on C23 of OTUB1. OTUB1 (10 μg) recombinant protein was incubated with EN523 (50 μM) for 30 min, after which the protein was precipitated and digested with trypsin and tryptic digests were analyzed by LC-MS/MS to identify modified sites. **(e)** OTUB1 DUB activity monitored by cleavage of K48 diubiquitin. Recombinant OTUB1 were pre-incubated with DMSO or EN523 (50 μM) for 1 h. After pre-incubation, OTUB1 was added to a mixture of diubiquitin and UBE2D1. The appearance of mono-Ub was monitored by Western blotting. **(f)** Structure of alkyne-functionalized EN523 probe—NJH-2-075. **(g)** Gel-based ABPP of NJH-2-075. Vehicle DMSO or NJH-2-075 were pre-incubated with OTUB1 for 30 min at 37 °C prior to IA-rhodamine labeling (500 nM, 30 min room temperature). OTUB1 was then separated by SDS/PAGE and in-gel fluorescence was assessed. Also shown is silver staining showing protein loading. **(h)** NJH-2-075 labeling of recombinant OTUB1. OTUB1 (0.5 μg) was labeled with DMSO or NJH-2-075 for 1.5 h at 37° C, after which rhodamine-azide was appended by CuAAC, OTUB1 was separated by SDS/PAGE and in-gel fluorescence was assessed. Also shown is silver staining showing protein loading. **(i)** NJH-2-075 engagement of OTUB1 in HEK293T cells. HEK293T cells were treated with DMSO vehicle or NJH-2-075 (50 μM) for 2 h, after which cell lysates were subjected to CuAAC with biotin picolyl azide and NJH-2-075 labeled proteins were subjected to avidin pulldown, eluted, separated by SDS/PAGE, and blotted for OTUB1 and vinculin. Both input lysate and pulldown levels are shown. Gels or blots shown in **(c, e, g, h, i)** are representative of n=3 biologically independent samples/group.

An alkyne-functionalized probe of EN523—NJH-2-075—was then synthesized with the goal of assessing whether this ligand is able to engage OTUB1 in cells **(Fig. 2f)**. NJH-2-075 retained binding to OTUB1 *in vitro*, as shown by: 1) gel-based ABPP demonstrating competition of NJH-2-075 against IA-rhodamine labeling of recombinant OTUB1 and; 2) direct labeling of recombinant OTUB1 by NJH-2-075 visualized by copper-catalyzed azide-alkyne cycloaddition (CuAAC) of azide-functionalized rhodamine to NJH-2-075-labeled OTUB1 and monitoring in-gel fluorescence **(Fig 2g-2h).** We demonstrated NJH-2-075 engagement of OTUB1 in cells by enrichment of endogenous OTUB1 through NJH-2-075 as compared with vehicle treatment in HEK293T cells, *ex situ* CuAAC of azide-functionalized biotin to NJH-2-075-labeled proteins and subsequent avidin-enrichment and blotting for OTUB1 **(Fig. 2i)**. We further showed that an unrelated protein vinculin was not enriched by NJH-2-075 **(Fig. 2i)**. Collectively, these data highlighted EN523 as a promising covalent OTUB1 ligand that targets a non-catalytic and allosteric C23 on OTUB1 without inhibiting OTUB1 deubiquitination activity and is able to engage OTUB1 in cells.

To demonstrate the feasibility of using EN523 as an OTUB1 recruiting module of a heterobifunctional DUBTAC, we turned to the mutant ΔF508-CFTR chloride channel as a proof-of-concept case where protein stabilization would be therapeutically desirable. ΔF508, a frameshift mutation caused by deletion at codon 508 in exon 10 of CFTR, resulting in the absence of a phenylalanine residue, is the most common mutation which induces the Cystic Fibrosis phenotype ^21^. This mutation causes the protein to become unstable and actively ubiquitinated with K48 ubiquitin chains, resulting in degradation prior to trafficking from the endoplasmic reticulum to the cell surface ^13,21^. Previous studies have shown the feasibility of stabilizing mutant CFTR not only by genetic and pharmacological inhibition of the cognate E3 ligase RNF5 but recently also through targeted recruitment of DUBs using a genetically encoded and engineered DUB targeted to CFTR through a CFTR-targeting nanobody ^22–24^. Importantly for our work, suitable CFTR-targeting small-molecule ligands exist in Lumacaftor, a drug for cystic fibrosis developed by Vertex, which acts as a chemical chaperone for ΔF508-CFTR to help stabilize its folding, leading to increased trafficking of ΔF508-CFTR to the cell membrane and partial restoration of function ^25^. Despite this chaperoning activity, the vast majority of ΔF508-CFTR is still actively ubiquitinated and degraded, making the potential of a synergistic effect via DUBTAC-induced deubiquitination a therapeutically attractive option. With this in mind, we synthesized DUBTACs linking the OTUB1 recruiter EN523 to the CFTR chaperone lumacaftor with two different C3 or C5 alkyl linkers—NJH-2-056 and NJH-2-057 **(Figure 3a-3b)**. We confirmed that these two DUBTACs still engaged recombinant OTUB1 *in vitro* by gel-based ABPP **(Figure 3c-3d).** The DUBTACs were then tested in CFBE41o-4.7 human bronchial epithelial cells expressing ΔF508-CFTR, a human cystic fibrosis bronchial epithelial cell line, alongside lumacaftor or EN523 treatment alone. While neither NJH-2-056, lumacaftor, nor EN523 treatment altered mutant CFTR levels, we observed a robust and significant increase in CFTR protein levels with NJH-2-057 treatment **(Figure 3e-3f).** This stabilization was dose-responsive, with clear stabilization occurring with 8 μM, and time-dependent with stabilization at 10 μM evident starting at 16 h **(Extended Data Figure 2).** We further confirmed that the stabilized protein was CFTR using three additional commercially available CFTR antibodies **(Extended Data Figure 3)** and showed that the DUBTAC-stabilized CFTR band was attenuated upon CFTR knockdown **(Extended Data Figure 4).** The CFTR smear that we observed in the blot with NJH-2-057 treatment is consistent with previous studies investigating CFTR and are also consistent with what we observed with treatment of cells with the proteasome inhibitor bortezomib, and likely represents a combination of differential glycosylation states, other forms of ubiquitination on CFTR that may not be removed by OTUB1 (e.g. K63 ubiquitin chains), and previously observed anomalous migration of CFTR on SDS/PAGE due to the presence of SDS-resistant ternary structures within the protein **(Extended Data Figure 5)**^21,26^. Based on the molecular weight of the darkest part of the CFTR blot >225 kDa, we conjecture that we are stabilizing the fully mature glycosylated form of mutant CFTR **(Figure 3e).**

**Figure 3.**
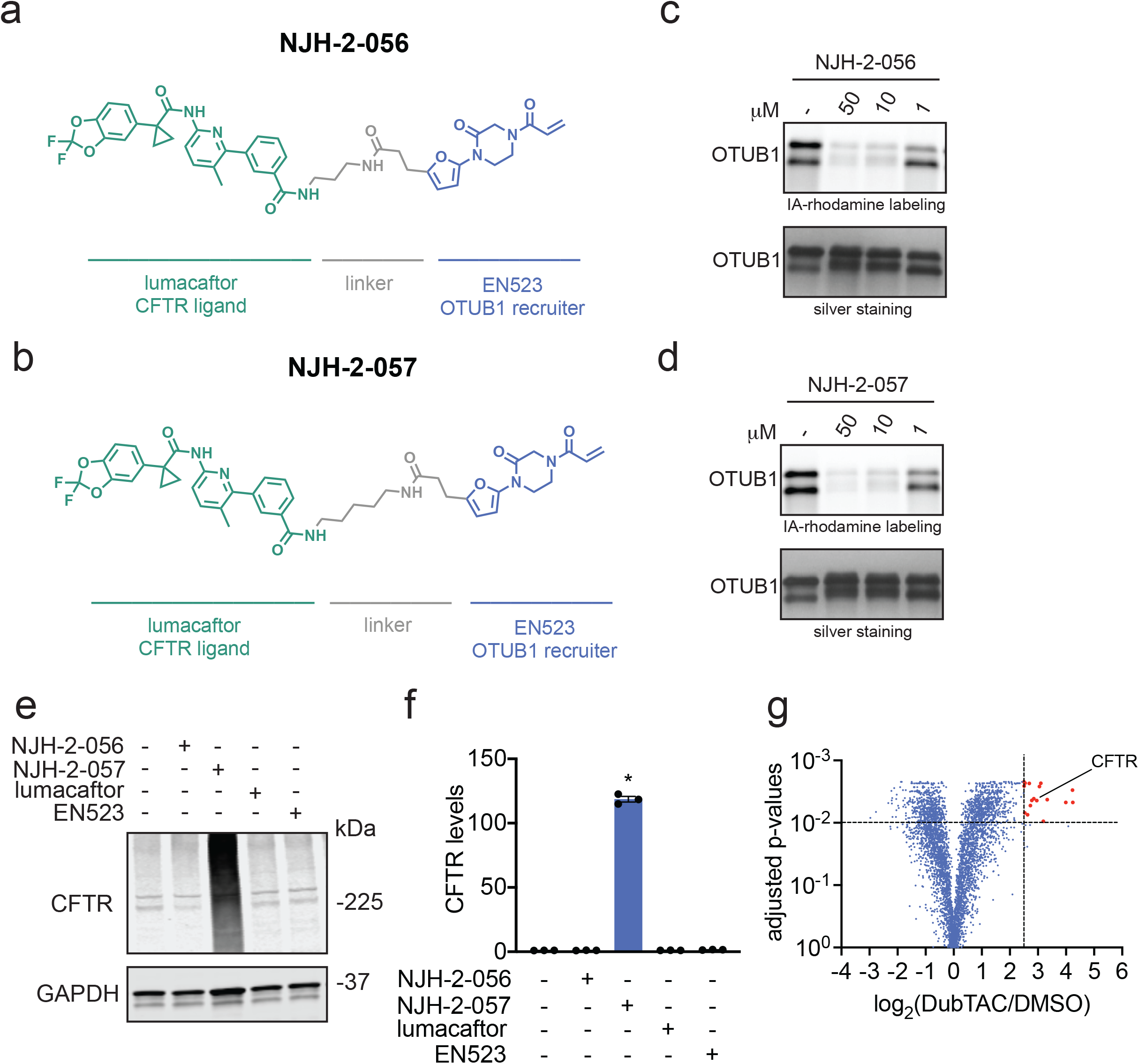
DUBTAC against mutant CFTR. **(a, b)** Structures of NJH-2-056 and NJH-2-057; these DUBTACs against mutant CFTR protein are formed by linking CFTR ligand lumacaftor to OTUB1 recruiter EN523 through C3 and C5 alkyl linkers, respectively. **(c, d)** Gel-based ABPP analysis of NJH-2-056 and NJH-2-057 against OTUB1. Vehicle DMSO or DUBTACs were preincubated with recombinant OTUB1 for 30 min at 37 °C prior to addition of IA-rhodamine (100 nM) for 30 min at room temperature. OTUB1 was run on SDS/PAGE and in-gel fluorescence was assessed. Protein loading was assessed by silver staining. **(e)** Effect of DUBTACs on mutant CFTR levels. CFBE41o-4.7 cells expressing ΔF508-CFTR were treated with vehicle DMSO, NJH-2-056 (10 μM), NJH-2-057 (10 μM), lumacaftor (10 μM), or EN523 (10 μM) for 24 h, and mutant CFTR and loading control GAPDH levels were assessed by Western blotting. **(f)** Quantification of the experiment described in **(e). (g)** TMT-based quantitative proteomic profiling of NJH-2-057 treatment. CFBE41o-4.7 cells expressing ΔF508-CFTR were treated with vehicle DMSO or NJH-2-057 (10 μM) for 24 h. Data shown are from n=3 biologically independent samples/group. Full data for this experiment can be found in **Table S3.** Gels shown in **(c, d, e)** are representative of n=3 biologically independent samples/group. Data in **(f)** show individual biological replicate values and average ± sem from n=3 biologically independent samples/group. Statistical significance was calculated with unpaired two-tailed Student’s t-tests in **(f)** and is expressed as *p<0.05.

To further validate our Western blotting data for CFTR stabilization and to assess the proteome-wide activity of NJH-2-057, we performed a TMT-based quantitative proteomic analysis of NJH-2-057 treated cells. Satisfyingly, the proteomic analysis showed CFTR amongst the most robustly stabilized proteins (ratio of 7.8 comparing NJH-2-057 to vehicle treatment, **Figure 3g; Table S4**). While there were additional proteins with significant changes in abundance levels, we only observed 21 proteins that were significantly stabilized by >5-fold with an adjusted p-value <0.01 compared to vehicle-treated controls out of 4552 total quantified proteins **(Figure 3g; Table S4)**. These additionally observed changes in protein abundance levels by the DUBTAC may represent either EN523 or Lumacaftor-specific changes or may be indication that we may be disrupting endogenous OTUB1 function for proteins showing decreased abundance levels.

To further confirm that the robust stabilization in mutant CFTR levels conferred by NJH-2-057 was due to the proposed on-target activity, we demonstrated that the stabilization of CFTR was attenuated by lumacaftor or EN523 pre-treatment, indicating that the stabilization by the DUBTAC was due to targets engaged by both lumacaftor and EN523 **(Figure 4a-4b).** Further verifying that the CFTR stabilization was dependent on OTUB1, OTUB1 knockdown significantly attenuated mutant CFTR stabilization by NJH-2-057 **(Figure 4c-4d)**.

**Figure 4.**
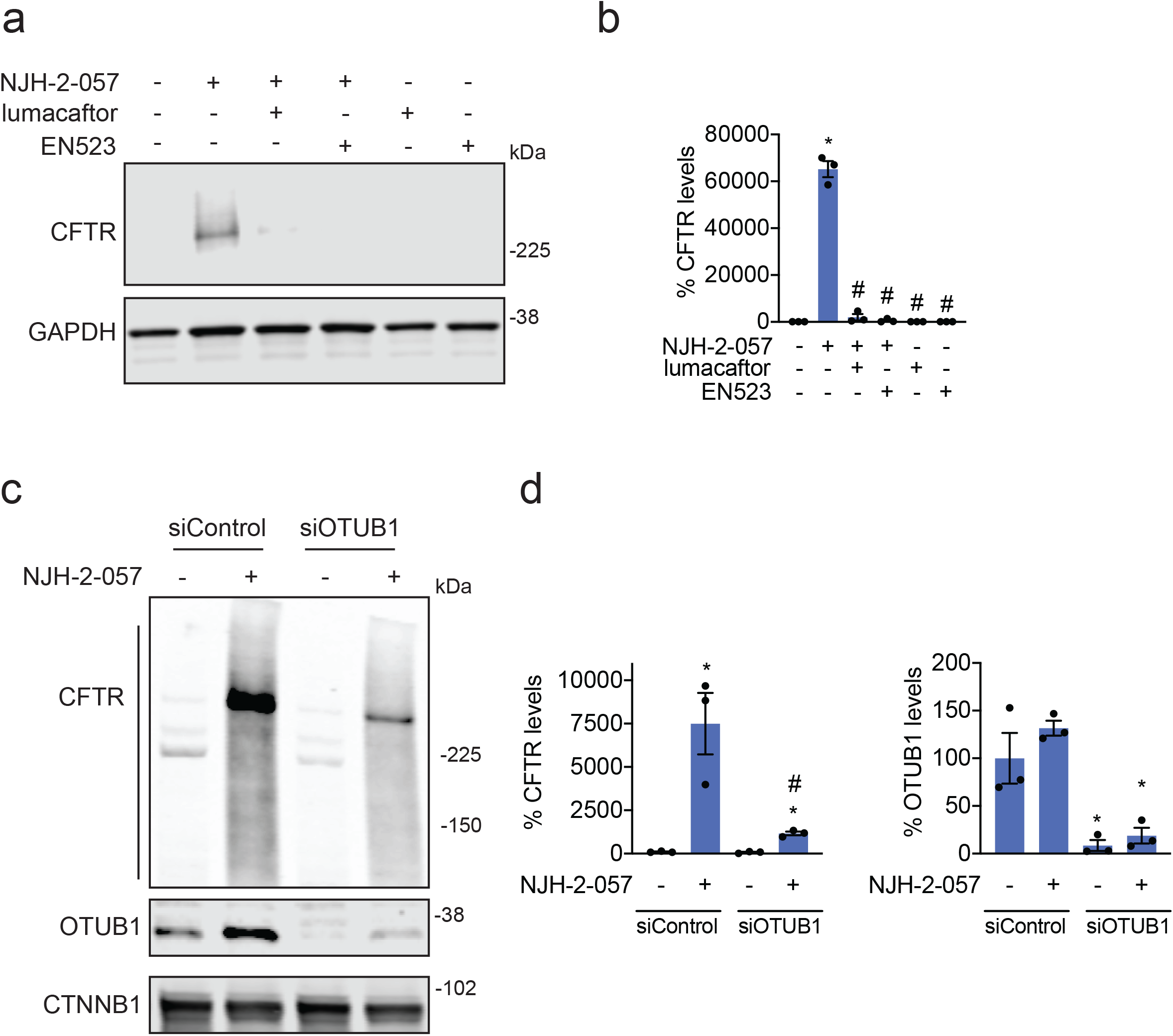
Characterizing the mechanism of the CFTR DUBTAC NJH-2-057. **(a)** Effect of lumacaftor or EN523 pre-incubation on NJH-2-057 DUBTAC-mediated stabilization of mutant CFTR levels. CFBE41o-4.7 cells expressing ΔF508-CFTR were pre-treated with vehicle DMSO, lumacaftor (100 μM), or EN523 (100 μM) for 1 h prior to treatment with NJH-2-057 (10 μM) for 24 h. Mutant CFTR and loading control GAPDH levels were assessed by Western blotting. **(b)** Quantification of the experiment described in **(a). (c)** Effect of OTUB1 knockdown on NJH-2-057 DUBTAC-mediated mutant CFTR stabilization. CFBE41o-4.7 cells expressing ΔF508-CFTR were transiently transfected with siControl or siOTUB1 oligonucleotides for 48 h prior to treatment of cells with vehicle DMSO or NJH-2-057 (10 μM) for 16 h. Mutant CFTR, OTUB1, and loading control GAPDH levels were assessed by Western blotting. **(d)** Levels of mutant CFTR and OTUB1 from the experiment described in **(c)**. Gels shown in **(a, c)** are representative of n=3 biologically independent samples/group. Data in **(b, d)** show individual biological replicate values and average ± sem from n=3 biologically independent samples/group. Statistical significance in was calculated with unpaired two-tailed Student’s t-tests in **(b, d)** and is expressed as *p<0.05 compared to vehicle-treated control in **(b)** and vehicle-treated siControl in **(d)** and #p<0.05 compared to the NJH-2-057 treated group in **(b)** and NJH-2-057 treated siControl group for CFTR levels in **(d)**.

Finally, we performed a competitive isoTOP-ABPP study to assess the overall proteome-wide selectivity and cysteine-reactivity of NJH-2-057 treatment in CFBE41o-4.7 cells expressing ΔF508-CFTR **(Extended Data Figure 6; Table S3)**. Out of 1270 IA-alkyne probe-modified peptides quantified in two out of three biological replicates, there were only 5 targets that showed isotopically light to heavy or control to NJH-2-057 treatment probe-modified peptide ratios of >4 with an adjusted p-value <0.05: VDAC2 C76, TUBB1 C201, RLF C744, and VDAC2 C47, and VDAC3 C66. None of these targets are part of the ubiquitin-proteasome system and as far as we can tell at this point unlikely to influence the activity of our DUBTAC **(Extended Data Figure 6; Table S3)**. OTUB1 C23 was captured in our isoTOP-ABPP experiment, but only showed a ratio of 1.6 which would correspond to ~60% target occupancy **(Table S3)**. This likely indicates that the observed CFTR stabilization by NJH-2-057 is occurring through relatively low levels of OTUB1 occupancy in cells. This is consistent with previous studies using covalent E3 ligase recruiters for targeted protein degradation applications showing that relatively minimal target occupancy of E3 ligases can still lead to robust degradation of target proteins due to the catalytic mechanism of action of the E3 ligases ^27–30^. We conjectured that a similar catalytic effect in a DUBTAC also leads to robust stabilization of the target protein with incomplete OTUB1 occupancy.

In this study, we have discovered a covalent small-molecule recruiter EN523 for the K48-ubiquitin chain-specific DUB OTUB1 and have shown proof-of-concept for targeted protein stabilization by small-molecule induced recruitment of a deubiquitinase to mutant CFTR as a protein of interest via a fully synthetic DUBTAC. While we showed early validation of the DUBTAC platform here, there are many avenues for future exploration. These include further optimization of DUB recruiters against OTUB1 to improve their potency and proteome-wide selectivity, as well as the discovery of new recruiters against other candidate DUBs. For exploring optimization of CFTR DUBTACs, further improvement of the linker between lumacaftor and the DUB recruiter could improve potency and degree of CFTR stabilization. In addition, elucidating the mechanism, structural underpinnings, and kinetics in the formation of the ternary complex formed between CFTR and OTUB1 and understanding how CFTR is deubiquitinated by the DUBTAC will be important. From a therapeutic point of view, it will be crucial to demonstrate that the lumacaftor-based DUBTAC results in increased membrane trafficking of CFTR protein and ultimately an increased functional capacity at the cellular level. Furthermore, better understanding of whether we are disrupting endogenous OTUB1 function would be important to understanding the mechanism and safety of DUBTACs.

Nonetheless, given our initial proof-of-concept for CFTR stabilization with a DUBTAC, there are many promising areas that could benefit from targeted deubiquitination of actively ubiquitinated and degraded proteins to provide therapeutic benefit. Targets that could benefit from a DUBTAC that already possess protein-targeting ligands include stabilizing BAX levels in the mitochondria to induce apoptosis, stabilizing STING for immunooncology applications, or stabilizing glucokinase in pancreatic cells for Type II diabetes ^31–34^. Other targets that would benefit from a DUBTAC would be various tumor suppressors such as TP53, CDKN1A, CDKN1C, AXIN1, PTEN, and others that are actively ubiquitinated and degraded to maintain cancer cell proliferation ^35^. One can also envision DUBTAC platforms for other types of ubiquitin chains with roles beyond degradation such as signaling, protein localization, and modulation of protein-protein interactions. Overall, our study puts forth the discovery of DUB recruiters and shows proof-of-concept for the DUBTAC platform for targeted protein stabilization via small molecule-induced deubiquitinase recruitment. In addition, our study underscores once more the utility of using chemoproteomics-enabled covalent ligand discovery platforms to facilitate development of unique induced proximity-based therapeutic modalities beyond targeted protein degradation.

## Acknowledgement

We thank the members of the Nomura Research Group and Novartis Institutes for BioMedical Research for critical reading of the manuscript. This work was supported by Novartis Institutes for BioMedical Research and the Novartis-Berkeley Center for Proteomics and Chemistry Technologies (NB-CPACT) for all listed authors. This work was also supported by the Nomura Research Group and the Mark Foundation for Cancer Research ASPIRE Award for DKN, NJH, LB, JNS, CCW, and BB. This work was also supported by grants from the National Institutes of Health (R01CA240981 for DKN) and the National Science Foundation Graduate Fellowship (for LB). We also thank Drs. Hasan Celik, Alicia Lund, and UC Berkeley’s NMR facility in the College of Chemistry (CoC-NMR) for spectroscopic assistance. Instruments in the CoC-NMR are supported in part by NIH S10OD024998.

## Author Contributions

NJH, LB, CCW, JNS, DKN conceived of the project idea, designed experiments, performed experiments, analyzed and interpreted the data, and wrote the paper. BB, SMB, DD performed experiments, analyzed and interpreted data, and provided intellectual contributions. MH contributed key critical intellectual contributions that enabled the DUBTAC technology. LMM, JMK, MS, JAT provided intellectual contributions to the project and overall design of the project.

---

**Table S1. Chemoproteomic analysis of DUBs.** Table of all probe-modified cysteines identified for 65 DUBs from 455 distinct isoTOP-ABPP experiments. This data is from our research group’s aggregate chemoproteomic experiments from various human cell lines wherein cell lysates were labeled with an alkyne-functionalized iodoacetamide (IA-alkyne) probe and taken through the isoTOP-ABPP procedure. Shown are the DUBs, probe-modified peptides, the site of the modified cysteine, the aggregate spectral counts for each particular probe-modified site identified across 455 experiments, and the total numbers of experiments for which the probe-modified peptide was observed.

**Table S2. Structures of covalent ligands screened against OTUB1.** Shown in **Tabs 1** and **2** are the structures of all compounds screened against OTUB1 using gel-based ABPP. Covalent ligand screen of cysteine-reactive libraries competed against IA-rhodamine labeling of recombinant OTUB1 to identify binders to OTUB1 by gel-based ABPP. Vehicle DMSO or cysteine-reactive covalent ligands (50 μM) were pre-incubated with OTUB1 for 30 min at room temperature prior to IA-rhodamine labeling (500 nM, 30 min room temperature). OTUB1 was then separated by SDS/PAGE and in-gel fluorescence was assessed and quantified. Gel-based ABPP data is shown in **Extended Data Figure 1a**. Quantification of gel-based ABPP data normalized to average of DMSO vehicle-treated controls in each gel is shown in **Tab 3**.

**Table S3. IsoTOP-ABPP analysis of NJH-2-057.** CFBE41o-4.7 cells expressing ΔF508-CFTR were treated with vehicle DMSO or NJH-2-057 for 8 h. Resulting cell lysates were labeled with IA-alkyne (200 μM) for 1 h and taken through the isoTOP-ABPP procedure Shown in red are the probe-modified peptides that showed isotopically light/heavy or control/NJH-2-57 ratios >4 with adjusted p-values <0.05. The data are from n=3 biologically independent samples/group.

**Table S4. TMT-based quantitative proteomic profiling of NJH-2-057 treatment.** CFBE41o-4.7 cells expressing ΔF508-CFTR were treated with vehicle DMSO or NJH-2-057 (10 μM) for 24 h. Data shown are from n=3 biologically independent samples/group.

## Online Methods

### Materials

Cysteine-reactive covalent ligand libraries were either previously synthesized and described or for the compounds starting with “EN” were purchased from Enamine, including EN523. Lumacaftor was purchased from Medchemexpress LLC.

### Cell Culture

CFBE41o-4.7 ΔF508-CFTR Human CF Bronchial Epithelial cells were purchased from Millipore Sigma (SCC159). CFBE41o-4.7 ΔF508-CFTR Human CF Bronchial Epithelial cells were cultured in MEM (Gibco) containing 10% (v/v) fetal bovine serum (FBS) and maintained at 37 °C with 5% CO_2_.

### Gel-Based ABPP

Recombinant OTUB1 (0.1μg/sample) was pre-treated with either DMSO vehicle or covalent ligand or DUBTACs at 37 °C for 30 min in 25 μL of PBS, and subsequently treated with of IA-Rhodamine (concentrations designated in figure legends) (Setareh Biotech) at room temperature for 1 h. The reaction was stopped by addition of 4×reducing Laemmli SDS sample loading buffer (Alfa Aesar). After boiling at 95 °C for 5 min, the samples were separated on precast 4−20% Criterion TGX gels (Bio-Rad). Probe-labeled proteins were analyzed by in-gel fluorescence using a ChemiDoc MP (Bio-Rad).

### NJH-2-057 Probe Labeling of Recombinant OTUB1

Recombinant and pure OTUB1 protein (0.5 μg) per sample per replicate was suspended in 50 μL total PBS. 1 μL of either DMSO or NJH-2-075 (to give final concentrations of 50, 10, 1, and 0.1 μM) was added, followed by a 1.5 h incubation at 37 °C. Next, 7.8 μL of a solution composed of 9.4 μL of 5mM Azide-Fluor 545 (in DMSO), 112 μL of TBTA ligand (Stock 1.7 mM in 4 parts t-butanol + 1 part DMSO), 37.5 μL of 50 mM TCEP (in water), and 37.5 μL of 50 mM Copper (II) sulfate was added to each sample and the samples were incubated for 1 hour at room temperature. Following CuAAC, 30 μL of Laemmli Sample Buffer (4 x) was added to each sample, vortexed and boiled for 6 min at 95 °C. Samples were loaded on an SDS/PAGE gel and analyzed for in-gel fluorescence.

### Deubiquitinase Activity Assay

Previously described methods were used to assess EN523 effects on OTUB1 activity ^20^. Recombinant OTUB1 (500 nM) was pre-incubated with DMSO or EN523 (50 μM) for 1 hr. To initiate assay pre-treated OTUB1 enzyme was mixed 1:1 with di-Ub reaction mix for final concentrations of 250 nM OTUB1, 1.5 µM di-Ub, 12.5 µM UBE2D1 and 5 mM DTT. The appearance of mono-Ub was monitored by Western blotting over time by removing a portion of the reaction mix and adding Laemmli’s buffer to terminaye reaction. Blot shown is a representative gel from n=3 biologically independent experiments/group.

### Labeling of Endogenous OTUB1 in HEK293T Cells with NJH-2-075 Probe

One plate of 70% confluent HEK293T cells per condition per replicate were treated with either DMSO vehicle or NJH-02-075 (50 μM) for 2 hours. Cells were harvested by scraping, suspended in 600 μL of PBS, lysed by probe sonication, and centrifuged for 10 min at 5000 rpm to remove debris. Lysate was normalized to 3.1 mg/mL and 85 μL removed for Western blotting analysis of input. 500 μL of lysate was then incubated for 1 hour at room temperature with 10 μL of 5 mM biotin picolyl azide (in water), 10 μL of 50mM TCEP (in water), 30 μL TBTA ligand (Stock 1.7 mM in 4 parts t-butanol + 1 part DMSO), and 10 μL of 50 mM Copper (II) sulfate. Following CuAAC, precipitated proteins were washed 3 x with cold methanol and resolubilized in 200 μL 1.2% SDS/PBS. To ensure solubility, proteins were heated to 90 °C for 5 min following resuspension. 1 mL of PBS was then added to each sample, followed by 50 μL of high-capacity streptavidin beads. Samples were then incubated overnight on a rocker at 4 °C. The following morning the samples were warmed to room temperature, and non-specific binding proteins were washed away with 3 x PBS washes followed by 3 x water washes. Beads were then resuspended in 100 μL PBS and 30 μL Laemmli Sample Buffer (4 x) and boiled for 13 min at 95 °C. Samples were vortexed and loaded onto an SDS/PAGE gel along with saved input samples for Western blotting analysis.

### Western Blotting

Proteins were resolved by SDS/PAGE and transferred to nitrocellulose membranes using the Trans-Blot Turbo transfer system (Bio-Rad). Membranes were blocked with 5% BSA in Tris-buffered saline containing Tween 20 (TBS-T) solution for 30 min at RT, washed in TBS-T, and probed with primary antibody diluted in recommended diluent per manufacturer overnight at 4°C. After 3 washes with TBS-T, the membranes were incubated in the dark with IR680- or IR800-conjugated secondary antibodies at 1:10,000 dilution in 5 % BSA in TBS-T at RT for 1 h. After 3 additional washes with TBST, blots were visualized using an Odyssey Li-Cor fluorescent scanner. The membranes were stripped using ReBlot Plus Strong Antibody Stripping Solution (EMD Millipore) when additional primary antibody incubations were performed. Antibodies used in this study were CFTR (Cell Signaling Technologies, Rb mAb #78335, **Figures 3 and 4**), CFTR (R&D Systems, Ms mAb, #MAB25031, **Extended Data Figure 3**), CFTR (Millipore, Ms mAb, #MAB3484, **Extended Data Figure 3**), CFTR (Prestige, Rb pAb, #HPA021939, **Extended Data Figure 3**), GAPDH (Proteintech, Ms mAb, #60004-1-Ig), OTUB1 (Abcam, Rb mAb, #ab175200, [EPR13028(B)]), and CTNNB1 (Cell Signaling Technologies, Rb mAb, #8480).

### IsoTOP-ABPP Chemoproteomic Experiments

IsoTOP-ABPP studies were done as previously reported ^14,27,36^. Our aggregate chemoproteomic data analysis of DUBs were obtained from 455 distinct isoTOP-ABPP experiments performed in the Nomura Research Group. These data are aggregated from various human cell lines, including 231MFP, A549, HeLa, HEK293T, HEK293A, UM-Chor1, PaCa2, PC3, HUH7, NCI-H460, THP1, SKOV3, U2OS, and K562 cells. Some of the chemoproteomic data have been previously reported as part of other studies ^30,36–45^. All of the isoTOP-ABPP datasets were prepared as described below using the IA-alkyne probe. Cells were lysed by probe sonication in PBS and protein concentrations were measured by BCA assay. Cells were treated for 4 h with either DMSO vehicle or EN4 (from 1,000x DMSO stock) before cell collection and lysis. Proteomes were subsequently labeled with IA-alkyne labeling (100 μM for DUB ligandability analysis and 200 μM for profiling cysteine-reactivity of NJH-2-057) for 1 h at room temperature. CuAAC was used by sequential addition of tris(2-carboxyethyl)phosphine (1 mM, Strem, 15-7400), tris[(1-benzyl-1H-1,2,3-triazol-4-yl)methyl]amine (34 μM, Sigma, 678937), copper(II) sulfate (1 mM, Sigma, 451657) and biotin-linker-azide—the linker functionalized with a tobacco etch virus (TEV) protease recognition sequence as well as an isotopically light or heavy valine for treatment of control or treated proteome, respectively. After CuAAC, proteomes were precipitated by centrifugation at 6,500*g*, washed in ice-cold methanol, combined in a 1:1 control:treated ratio, washed again, then denatured and resolubilized by heating in 1.2% SDS–PBS to 80 °C for 5 min. Insoluble components were precipitated by centrifugation at 6,500*g* and soluble proteome was diluted in 5 ml 0.2% SDS–PBS. Labeled proteins were bound to streptavidin-agarose beads (170 μl resuspended beads per sample, Thermo Fisher, 20349) while rotating overnight at 4 °C. Bead-linked proteins were enriched by washing three times each in PBS and water, then resuspended in 6 M urea/PBS, and reduced in TCEP (1 mM, Strem, 15-7400), alkylated with iodoacetamide (18 mM, Sigma), before being washed and resuspended in 2 M urea/PBS and trypsinized overnight with 0.5 μg /μL sequencing grade trypsin (Promega, V5111). Tryptic peptides were eluted off. Beads were washed three times each in PBS and water, washed in TEV buffer solution (water, TEV buffer, 100 μM dithiothreitol) and resuspended in buffer with Ac-TEV protease (Invitrogen, 12575-015) and incubated overnight. Peptides were diluted in water and acidified with formic acid (1.2 M, Fisher, A117-50) and prepared for analysis.

### IsoTOP-ABPP Mass Spectrometry Analysis

Peptides from all chemoproteomic experiments were pressure-loaded onto a 250 μm inner diameter fused silica capillary tubing packed with 4 cm of Aqua C18 reverse-phase resin (Phenomenex, 04A-4299), which was previously equilibrated on an Agilent 600 series high-performance liquid chromatograph using the gradient from 100% buffer A to 100% buffer B over 10 min, followed by a 5 min wash with 100% buffer B and a 5 min wash with 100% buffer A. The samples were then attached using a MicroTee PEEK 360 μm fitting (Thermo Fisher Scientific p-888) to a 13 cm laser pulled column packed with 10 cm Aqua C18 reverse-phase resin and 3 cm of strong-cation exchange resin for isoTOP-ABPP studies. Samples were analyzed using an Q Exactive Plus mass spectrometer (Thermo Fisher Scientific) using a five-step Multidimensional Protein Identification Technology (MudPIT) program, using 0, 25, 50, 80 and 100% salt bumps of 500 mM aqueous ammonium acetate and using a gradient of 5–55% buffer B in buffer A (buffer A: 95:5 water:acetonitrile, 0.1% formic acid; buffer B 80:20 acetonitrile:water, 0.1% formic acid). Data were collected in data-dependent acquisition mode with dynamic exclusion enabled (60 s). One full mass spectrometry (MS1) scan (400–1,800 mass-to-charge ratio (*m/z*)) was followed by 15 MS2 scans of the *n*th most abundant ions. Heated capillary temperature was set to 200 °C and the nanospray voltage was set to 2.75 kV.

Data were extracted in the form of MS1 and MS2 files using Raw Extractor v.1.9.9.2 (Scripps Research Institute) and searched against the Uniprot human database using ProLuCID search methodology in IP2 v.3-v.5 (Integrated Proteomics Applications, Inc.) ^46^. Cysteine residues were searched with a static modification for carboxyaminomethylation (+57.02146) and up to two differential modifications for methionine oxidation and either the light or heavy TEV tags (+464.28596 or +470.29977, respectively). Peptides were required to be fully tryptic peptides and to contain the TEV modification. ProLUCID data were filtered through DTASelect to achieve a peptide false-positive rate below 5%. Only those probe-modified peptides that were evident across two out of three biological replicates were interpreted for their isotopic light to heavy ratios. For those probe-modified peptides that showed ratios greater than two, we only interpreted those targets that were present across all three biological replicates, were statistically significant and showed good quality MS1 peak shapes across all biological replicates. Light versus heavy isotopic probe-modified peptide ratios are calculated by taking the mean of the ratios of each replicate paired light versus heavy precursor abundance for all peptide-spectral matches associated with a peptide. The paired abundances were also used to calculate a paired sample *t*-test *P* value in an effort to estimate constancy in paired abundances and significance in change between treatment and control. *P* values were corrected using the Benjamini–Hochberg method.

### Knockdown studies

RNA interference was performed using siRNA purchased from Dharmacon. CFBE41o-4.7 cells were seeded at 400,000 cells per 6 cm plate and allowed to adhere overnight. Cells were transfected with 33 nM of either nontargeting (ON-TARGETplus Non-targeting Control Pool, Dharmacon #D-001810-10-20) or anti-CFTR siRNA (Dharmacon, custom) using 8 μL of transfection reagent: either DharmaFECT 1 (Dharmacon #T-2001-02), DharmaFECT 4 (Dharmacon, T-2004-02) or Lipofectamine 2000 (ThermoFisher #11668027). Transfection reagent was added to OPTIMEM (ThermoFisher #31985070) media, allowed to incubate for 5 min at room temperature. Meanwhile siRNA was added to an equal amount of OPTIMEM. Solutions of transfection reagent and siRNA in OPTIMEM were then combined and allowed to incubate for 30 minutes at room temperature. These combined solutions were diluted with complete MEM to provide 33nM siRNA and 8 μL of transfection reagent per 4 mL MEM, and the media exchanged. Cells were incubated with transfection reagents for 24h, at which point the media replaced with media containing DMSO or 10 μM NJH-2-057 and incubated for another 24h. Cells were then harvested, and protein abundance analyzed by Western blotting.

### Quantitative TMT Proteomics Analysis

Quantitative TMT-based proteomic analysis was performed as previously described ^27^. Acquired MS data was processed using Proteome Discoverer v. 2.4.0.305 software (Thermo) utilizing Mascot v 2.5.1 search engine (Matrix Science, London, UK) together with Percolator validation node for peptide-spectral match filtering ^47^. Data was searched against Uniprot protein database (canonical human sequences, EBI, Cambridge, UK) supplemented with sequences of common contaminants. Peptide search tolerances were set to 10 ppm for precursors, and 0.8 Da for fragments. Trypsin cleavage specificity (cleavage at K, R except if followed by P) allowed for up to 2 missed cleavages. Carbamidomethylation of cysteine was set as a fixed modification, methionine oxidation, and TMT-modification of N-termini and lysine residues were set as variable modifications. Data validation of peptide and protein identifications was done at the level of the complete dataset consisting of combined Mascot search results for all individual samples per experiment via the Percolator validation node in Proteome Discoverer. Reporter ion ratio calculations were performed using summed abundances with most confident centroid selected from 20 ppm window. Only peptide-to-spectrum matches that are unique assignments to a given identified protein within the total dataset are considered for protein quantitation. High confidence protein identifications were reported using a Percolator estimated <1% false discovery rate (FDR) cut-off. Differential abundance significance was estimated using ANOVA with Benjamini-Hochberg correction to determine adjusted p-values.

### Data Availability Statement

The datasets generated during and/or analyzed during the current study are available from the corresponding author on reasonable request.

### Code Availability Statement

Data processing and statistical analysis algorithms from our lab can be found on our lab’s Github site: https://github.com/NomuraRG, and we can make any further code from this study available at reasonable request.

### Chemical Synthesis and Characterization

Starting materials, reagents and solvents were purchased from commercial suppliers and were used without further purification unless otherwise noted. All reactions were monitored by TLC (TLC Silica gel 60 F_254_, Sepulco Millipore Sigma). Reaction products were purified by flash column chromatography using a Biotage Isolera with Biotage Sfar® or Silicycle normal-phase silica flash columns (5 g, 10 g, 25 g, or 40 g). 1H NMR and 13C NMR spectra were recorded on a 400 MHz Bruker Avance I spectrometer or a 600 MHz Bruker Avance III spectrometer equipped with a 5 mm 1H/BB Prodigy cryo-probe. Chemical shifts are reported in parts per million (ppm, δ) downfield from tetramethylsilane (TMS). Coupling constants (J) are reported in Hz. Spin multiplicities are described as br (broad), s (singlet), d (doublet), t (triplet), q (quartet) and m (multiplet).

### SYNTHESIS OF NJH-2-057

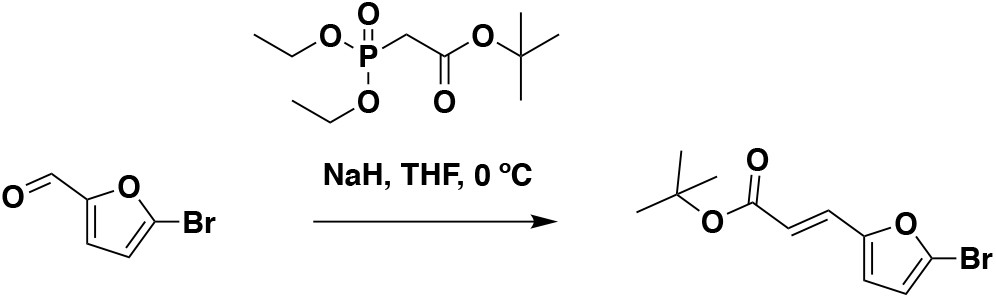

**tert-butyl (*E*)-3-(5-bromofuran-2-yl)acrylate:** tert-butyl diethylphosphonoacetate (971 mg, 0.908 mL, 3.85 mmol) was dissolved in THF (22 mL) and the solution cooled to 0 °C. Then, 2-bromofuran-2carbaldehyde (613 mg, 3.50 mmol) was added portion-wise over 5 minutes. The reaction was stirred for 20 minutes at 0 °C as a gummy solid precipitated and then water was added. The resulting mixture was extracted with EtOAc, combined organic extracts were washed with brine, dried over Na_2_SO_4_, concentrated, and purified by silica gel chromatography (0-15% EtOAc/Hex) to provide the title compound as a light yellow oil (782 mg, 2.86 mmol, 82%). 1H NMR (400 MHz, CDCl_3_) δ 7.26 (d, J = 15.7 Hz, 1H), 6.55 (d, J = 3.5 Hz, 1H), 6.42 (d, J = 3.4 Hz, 1H), 6.29 (d, J = 15.7 Hz, 1H), 1.55 (s, 9H).

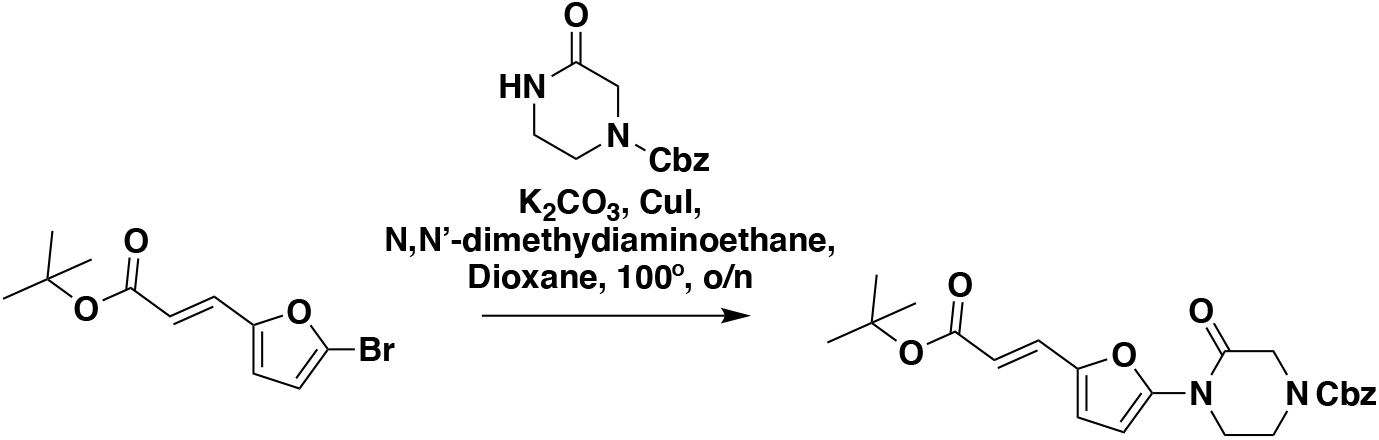

**benzyl (*E*)-4-(5-(3-(tert-butoxy)-3-oxoprop-1-en-1-yl)furan-2-yl)-3-oxopiperazine-1-carboxylate:** tert-butyl (*E*)-3-(5-bromofuran-2-yl)acrylate (1.62 g, 5.94 mmol) was dissolved in dioxane (30 mL) and benzyl 3-oxopiperazine-1-carboxylate (1.4 g, 5.94 mmol), K_2_CO_3_ (2.46 g, 17.8 mmol), *N,N*’-dimethyldiaminoethane (0.167 mL, 1.49 mmol), and CuI (114 mg, 0.59 mmol) were added. The mixture was stirred under nitrogen at reflux for 40 h, then cooled to rt. 5 mL saturated aq. NH_4_Cl was added and the mixture stirred for 30 min. Then the mixture was diluted in EtOAc, filtered through celite, water was added, the mixture partitioned, and the aqueous layer extracted with EtOAc. The extracts were combined, washed with brine, dried over Na_2_SO_4_, concentrated, and purified by silica gel chromatography (0-35% EtOAc/Hex) to provide the title compound as an orange oil (1.95 g, 4.59 mmol, 77%). LC/MS [M+2H-tBu]^+^ m/z calc. 371.18, found 373.1. 1H NMR (400 MHz, DMSO) δ 7.45 – 7.24 (m, 6H), 6.98 (s, 1H), 6.57 (s, 1H), 6.08 (dd, J = 15.7, 3.4 Hz, 1H), 5.14 (dd, J = 4.4, 2.3 Hz, 2H), 4.22 (s, 2H), 4.01 (s, 2H), 3.77 (s, 2H), 1.47 (s, 9H).

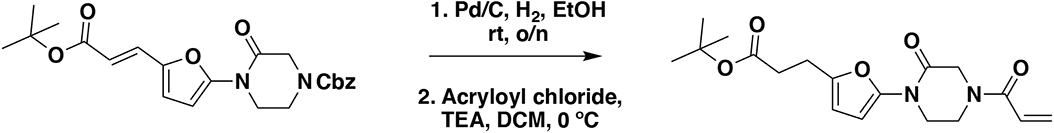

**tert-butyl 3-(5-(4-acryloyl-2-oxopiperazin-1-yl)furan-2-yl)propanoate (Intermediate 1):** benzyl (*E*)-4-(5-(3-(tert-butoxy)-3-oxoprop-1-en-1-yl)furan-2-yl)-3-oxopiperazine-1-carboxylate (1.95 g, 4.59 mmol) was dissolved in EtOH (25 mL) and Pd/C (200 mg, 10% wt. Pd) was added. The reaction was placed under an atmosphere of H_2_ and stirred vigorously overnight, before being filtered through celite twice and concentrated. The crude product was then redissolved in DCM (25 mL), cooled to 0 °C and treated with TEA (1.28 mL, 9.18 mmol) before a solution of acryloyl chloride (445 μL, 5.51 mmol) in DCM (5 mL) was added over 2 minutes. After stirring for 20 min, water was added and the mixture extracted with DCM three times. Combined organic extracts were washed with brine, dried over Na_2_SO_4_, concentrated, and the resulting crude oil was purified by silica gel chromatography (0-75% EtOAc/Hex) to obtain the title compound as a light yellow oil (846 mg, 2.43 mmol, 53% over two steps). LC/MS [M+2H-tBu]^+^ m/z calc. 293.1, found 293.1. 1H NMR (400 MHz, CDCl_3_) δ 6.64 – 6.46 (m, 1H), 6.41 (dd, J = 16.7, 2.0 Hz, 1H), 6.29 (d, J = 3.2 Hz, 1H), 6.04 (d, J = 3.3 Hz, 1H), 5.82 (dd, J = 10.2, 2.0 Hz, 1H), 4.42 (d, J = 24.9 Hz, 2H), 4.06 – 3.82 (m, 4H), 2.88 (t, J = 7.8 Hz, 2H), 2.54 (d, J = 7.6 Hz, 2H), 1.44 (s, 9H).

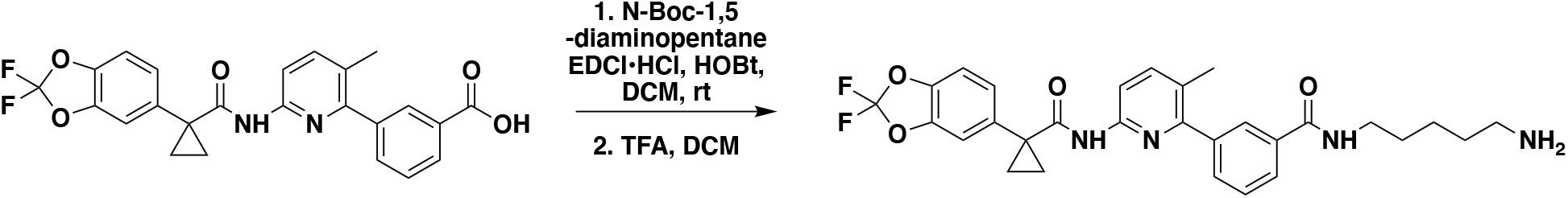

***N*-(5-aminopentyl)-3-(6-(1-(2,2-difluorobenzo[d][1,3]dioxol-5-yl)cyclopropane-1-carboxamido)-3-methylpyridin-2-yl)benzamide:** Lumacaftor (3-(6-(1-(2,2-difluorobenzo[d][1,3]dioxol-5-yl)cyclopropane-1-carboxamido)-3-methylpyridin-2-yl)benzoic acid) (181 mg, 0.40 mmol), tert-butyl (5-aminopentyl)carbamate (121 mg, 0.60 mmol), DIEA (350 μL, 2.00 mmol), and HOBt (54 mg, 0.4mmol) were dissolved in DCM (6 mL), followed by addition of EDCI HCl (153 mg, 0.50 mmol). The reaction was stirred at rt for 16 hours before water was added, the mixture partitioned, and the aqueous layer extracted with DCM twice. The combined organic extracts were washed with brine, dried over Na_2_SO_4_, concentrated, and the resulting crude oil was purified by silica gel chromatography (0-50% EtOAc/Hex) to obtain the title compound as a clear oil (240 mg, 0.38 mmol, 95%). LC/MS [M+H]^+^ m/z calc. 637.28, found 637.3. 1H NMR (400 MHz, CDCl_3_) δ 8.14 (d, J = 8.4 Hz, 1H), 7.84 (s, 1H), 7.80 (dt, J = 7.6, 1.6 Hz, 1H), 7.73 (s, 1H), 7.63 (d, J = 8.5 Hz, 1H), 7.57 (dt, J = 7.7, 1.5 Hz, 1H), 7.51 (t, J = 7.6 Hz, 1H), 7.27 (dd, J = 8.1, 1.8 Hz, 1H), 7.23 (d, J = 1.7 Hz, 1H), 7.12 (d, J = 8.2 Hz, 1H), 6.25 (s, 1H), 3.17 (d, J = 6.8 Hz, 2H), 4.61 (s, 1H), 3.49 (q, J = 7.0, 6.8, 6.3 Hz, 2H), 2.29 (s, 3H), 1.79 (q, J = 3.9 Hz, 2H), 1.56 (q, J = 7.2 Hz, 2H), 1.46 (s, 11H), 1.36 – 1.27 (m, 2H), 1.20 (q, J = 3.9 Hz, 2H), 0.97 – 0.89 (m, 2H). This Boc-protected amine (240 mg, 0.038 mmol) was dissolved in DCM (2 mL) and TFA (2 mL) was added and the solution stirred for 2 hours. The volatiles were then evaporated and the resulting oil redissolved in DCM and treated with aqueous saturated NaHCO_3_. The layers were separated and the aqueous layer was then extracted with DCM three times. The combined organic extracts were dried over Na_2_SO_4_, and concentrated to provide the title compound (184 mg, 0.34 mmol, 85 % over two steps) as a colorless oil which was used in the next step without further purification. LC/MS [M+H]^+^ m/z calc. 537.22, found 537.2. 1H NMR (400 MHz, CDCl_3_) δ 8.09 (d, J = 8.4 Hz, 1H), 7.80 (t, J = 1.8 Hz, 1H), 7.76 (dd, J = 7.7, 1.5 Hz, 1H), 7.69 (s, 1H), 7.59 (d, J = 8.5 Hz, 1H), 7.57 – 7.50 (m, 1H), 7.47 (t, J = 7.6 Hz, 1H), 7.23 (dd, J = 8.2, 1.7 Hz, 1H), 7.19 (d, J = 1.8 Hz, 1H), 7.08 (d, J = 8.2 Hz, 1H), 6.30 (s, 1H), 3.45 (q, J = 6.7 Hz, 2H), 2.74 (t, J = 6.8 Hz, 2H), 2.25 (s, 3H), 1.65 – 1.59 (m, 2H), 1.57 – 1.47 (m, 2H), 1.48 – 1.40 (m, 2H), 1.33 – 1.23 (m, 2H), 1.20 – 1.12 (m, 2H), 0.91 – 0.85 (m, 2H).

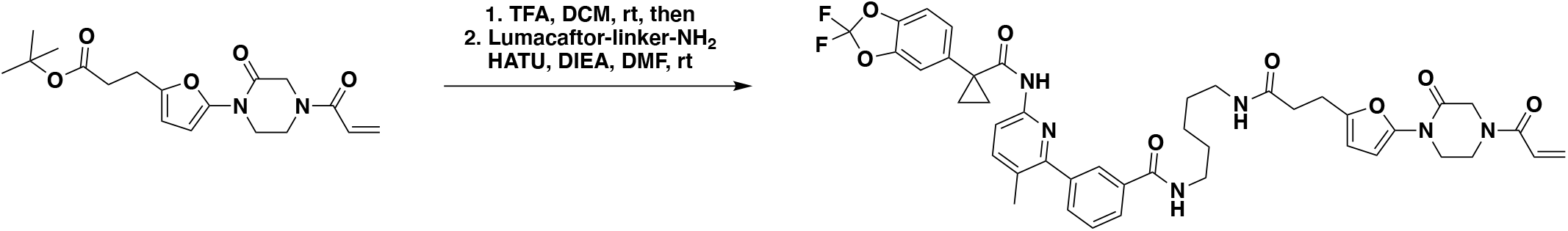

***N*-(5-(3-(5-(4-acryloyl-2-oxopiperazin-1-yl)furan-2-yl)propanamido)pentyl)-3-(6-(1-(2,2-difluorobenzo[d][1,3]dioxol-5-yl)cyclopropane-1-carboxamido)-3-methylpyridin-2-yl)benzamide (NJH-2-057):** Intermediate 1 (tert-butyl 3-(5-(4-acryloyl-2-oxopiperazin-1-yl)furan-2-yl)propanoate) (70 mg, 0.20 mmol) was dissolved in DCM (1.0 mL) and TFA (0.8 mL) was added and the solution stirred for 1 h until starting material was consumed as monitored by TLC. The volatiles were evaporated, DCM was added and evaporated again. The residue was dissolved in DMF (1.5 mL) and DIEA (150 μL, 0.86 mmol) was added followed by N-(5-aminopentyl)-3-(6-(1-(2,2-difluorobenzo[d][1,3]dioxol-5-yl)cyclopropane-1-carboxamido)-3-methylpyridin-2-yl)benzamide (54 mg, 0.1 mmol). HATU (152 mg, 0.4mmol) was then added and the mixture stirred for 1 h. Water was added, and the resulting suspension was extracted with DCM three times. Combined organic extracts were washed twice with 1M HCl twice, saturated NaHCO_3_, twice with 5% sat. LiCl, brine, and dried over Na_2_SO_4_ before being concentrated. The crude residue was purified by silica gel chromatography (0-4% MeOH/DCM) to obtain the title compound (35 mg, 0.043 mmol, 43%) as a white powder following lyopilization from 1:1 water:acetonitrile (2 mL). HRMS [M+H]^+^ m/z calc. 811.3262, found 811.3267. 1H NMR (600 MHz, CDCl_3_) δ 8.11 (d, J = 8.4 Hz, 1H), 7.85 (t, J = 1.8 Hz, 1H), 7.81 (dt, J = 7.8, 1.5 Hz, 1H), 7.71 (s, 1H), 7.61 (d, J = 8.5 Hz, 1H), 7.55 (dt, J = 7.7, 1.4 Hz, 1H), 7.49 (t, J = 7.6 Hz, 1H), 7.25 (dd, J = 8.2, 1.8 Hz, 1H), 7.21 (d, J = 1.8 Hz, 1H), 7.10 (d, J = 8.2 Hz, 1H), 6.53 (s, 1H), 6.41 (dd, J = 16.7, 1.8 Hz, 2H), 6.22 (d, J = 3.3 Hz, 1H), 6.03 (d, J = 3.3 Hz, 1H), 5.82 (dd, J = 10.4, 1.8 Hz, 2H), 4.54 – 4.32 (m, 2H), 4.07 – 3.79 (m, 4H), 3.45 (q, J = 6.6 Hz, 2H), 3.24 (q, J = 6.6 Hz, 2H), 2.91 (t, J = 7.3 Hz, 2H), 2.46 (t, J = 7.3 Hz, 2H), 2.27 (s, 3H), 1.77 (q, J = 3.9 Hz, 2H), 1.65 – 1.59 (m, 2H), 1.52 (p, J = 7.0 Hz, 2H), 1.40 – 1.32 (m, 2H), 1.18 (q, J = 3.9 Hz, 2H). 13C NMR (151 MHz, CDCl_3_) δ 171.7, 167.4, 165.0, 155.5, 148.9, 144.8, 144.1, 143.6, 141.0, 140.2, 134.9, 134.8, 131.8, 128.4, 127.5, 127.0, 126.6, 126.6, 126.3, 112.9, 112.4, 110.2, 107.4, 100.9, 39.7, 39.1, 31.2, 29.0, 24.2, 23.7, 19.2, 17.2.

### SYNTHESIS OF NJH-2-056

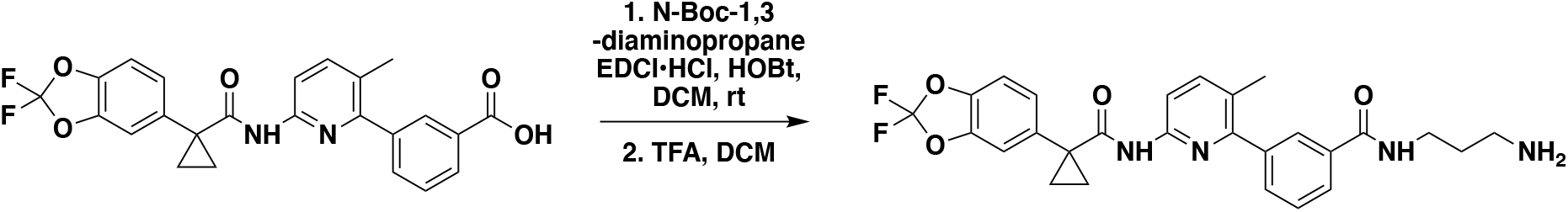

***N*-(3-aminopropyl)-3-(6-(1-(2,2-difluorobenzo[d][1,3]dioxol-5-yl)cyclopropane-1-carboxamido)-3-methylpyridin-2-yl)benzamide:** Lumacaftor (3-(6-(1-(2,2-difluorobenzo[d][1,3]dioxol-5-yl)cyclopropane-1-carboxamido)-3-methylpyridin-2-yl)benzoic acid) (18 mg, 0.04 mmol), tert-butyl (3-aminopropyl)carbamate (14 mg, 0.08 mmol), DIEA (35 μL, 0.20 mmol), and HOBt (5.4 mg, 0.04 mmol) were dissolved in DCM (1 mL), followed by the addition of EDCI HCl (15 mg, 0.05 mmol). The reaction was stirred at rt for 2 days before water was added, the mixture partitioned, and the aqueous layer extracted with DCM twice. The combined organic extracts were washed with brine, dried over Na_2_SO_4_, concentrated, and the resulting crude oil was purified by silica gel chromatography (0-60% EtOAc/Hex) to obtain the title compound as a clear oil (23 mg, 0.038 mmol, 94%). LC/MS [M+H]^+^ m/z calc. 609.24, found 609.3. 1H NMR (400 MHz, CDCl_3_) δ 8.13 (d, J = 8.4 Hz, 1H), 7.95 (s, 1H), 7.88 (d, J = 7.6 Hz, 1H), 7.74 (s, 1H), 7.62 (d, J = 8.5 Hz, 1H), 7.60 – 7.49 (m, 2H), 7.34 (s, 1H), 7.30 – 7.18 (m, 2H), 7.11 (d, J = 8.2 Hz, 1H), 4.96 (s, 1H), 3.54 (q, J = 6.2 Hz, 2H), 3.27 (q, J = 6.3 Hz, 2H), 2.31 (s, 3H), 1.78 (q, J = 3.9 Hz, 2H), 1.76 – 1.70 (m, 2H), 1.47 (s, 9H), 1.19 (q, J = 3.9 Hz, 2H). This Boc-protected amine (23 mg, 0.038 mmol) was dissolved in DCM (1 mL) and TFA (1 mL) was added and the solution stirred for 2 hours. The volatiles were then evaporated and the resulting oil redissolved in DCM and treated with aqueous saturated NaHCO_3_. The resulting mixture was then extracted with DCM three times, combined organic extracts dried over Na_2_SO_4_, concentrated to provide the title compound (15 mg, 0.029 mmol, 78%) as a colorless oil which was used in the next step without further purification. LC/MS [M+H]^+^ m/z calc. 509.19, found 509.2. 1H NMR (400 MHz, CDCl_3_) δ 10.73 (s, 1H), 8.96 (s, 1H), 8.66 (t, J = 5.7 Hz, 1H), 7.95 – 7.85 (m, 3H), 7.79 – 7.66 (m, 2H), 7.60 (d, J = 7.6 Hz, 1H), 7.56 – 7.49 (m, 2H), 7.41 – 7.30 (m, 2H), 3.33 (q, J = 6.4 Hz, 2H), 2.88 – 2.77 (m, 2H), 2.21 (s, 3H), 1.79 (p, J = 6.9 Hz, 2H), 1.52 (dd, J = 4.9, 2.5 Hz, 2H), 1.19 – 1.15 (m, 2H).

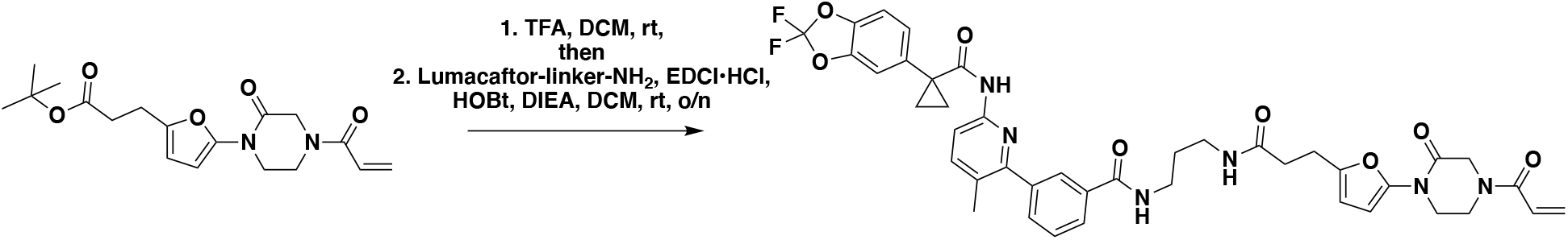

***N*-(3-(3-(5-(4-acryloyl-2-oxopiperazin-1-yl)furan-2-yl)propanamido)propyl)-3-(6-(1-(2,2-difluorobenzo[d][1,3]dioxol-5-yl)cyclopropane-1-carboxamido)-3-methylpyridin-2-yl)benzamide (NJH-2-056):** Intermediate 1 (tert-butyl 3-(5-(4-acryloyl-2-oxopiperazin-1-yl)furan-2-yl)propanoate) (14 mg, 0.04 mmol) was dissolved in DCM (0.6 mL) and TFA (0.3 mL) was added and the solution was stirred for 1 h at rt until starting material was consumed as monitored by TLC. Volatiles were evaporated, DCM was added and evaporated again. The residue was dissolved in DCM (1.5 mL) and DIEA (140 μL, 0.80 mmol) was added followed by *N*-(3-aminopropyl)-3-(6-(1-(2,2-difluorobenzo[d][1,3]dioxol-5-yl)cyclopropane-1-carboxamido)-3-methylpyridin-2-yl)benzamide (5.4 mg, 0.1 mmol). EDCI HCl (15 mg, 0.08 mmol) was then added and the mixture stirred for 16h. Water was added and the resulting suspension was extracted with DCM three times.

The combined organic extracts were washed with brine and dried over Na_2_SO_4_ before being concentrated. The crude residue was purified by silica gel chromatography (0-5% MeOH/DCM) to obtain the title compound (9.5 mg, 0.012 mmol, 30%) as a white powder following lyophilization from 1:1 water:acetonitrile (2 mL). HRMS [M+H]^+^ m/z calc. 783.2949, found 783.2954. 1H NMR (400 MHz, CDCl_3_) δ 8.09 (d, J = 8.4 Hz, 1H), 7.93 – 7.87 (m, 1H), 7.83 (dt, J = 7.5, 1.6 Hz, 1H), 7.72 (s, 1H), 7.59 (d, J = 8.5 Hz, 1H), 7.57 – 7.45 (m, 2H), 7.29 (s, 1H), 7.23 (dd, J = 8.2, 1.8 Hz, 1H), 7.19 (d, J = 1.7 Hz, 1H), 7.07 (d, J = 8.1 Hz, 1H), 6.50 (s, 1H), 6.43 – 6.33 (m, 2H), 6.19 (d, J = 3.2 Hz, 1H), 6.07 (d, J = 3.3 Hz, 1H), 5.81 (d, J = 10.1 Hz, 1H), 4.47 – 4.31 (m, 2H), 4.04 – 3.78 (m, 4H), 3.36 (q, J = 6.2 Hz, 2H), 3.32 – 3.23 (m, 2H), 2.96 (t, J = 7.2 Hz, 2H), 2.55 (t, J = 7.2 Hz, 2H), 2.26 (s, 3H), 1.74 (q, J = 3.9 Hz, 2H), 1.69 – 1.58 (m, 2H), 1.16 (q, J = 3.9 Hz, 2H). 13C NMR (151 MHz, CDCl_3_) δ 172.5, 171.8, 167.4, 165.0, 155.5, 149.8, 148.9, 145.0, 144.1, 143.6, 141.0, 140.2, 134.9, 134.6, 131.8, 131.7, 130.0, 128.5, 127.8, 127.0, 126.6, 126.5, 126.3, 112.9, 112.4, 110.2, 107.6, 101.3, 36.0, 35.9, 35.2, 31.2, 29.5, 24.4, 19.2, 17.2.

### SYNTHESIS OF NJH-2-075

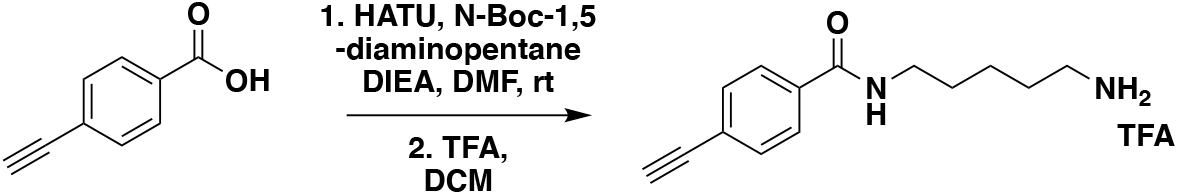

***N*-(5-aminopentyl)-4-ethynylbenzamide:** 4-ethynylbenzoic acid (27 mg, 0.19 mmol), N-Boc-1,5-diaminopentane (47 mg, 0.23 mmol), HOBt (26 mg, 0.19 mmol), and DIEA (165 μL, 0.95 mmol) were dissolved in DCM (1.5 mL) and EDCI-HCl (73 mg, 0.38 mmol) was added. After stirring the mixture for 16h at rt, water was added, the mixture partitioned, and the aqueous phase extracted with DCM. Combined organic extracts were washed with brine and dried over Na_2_SO_4_, concentrated, and the crude residue was purified by silica gel chromatography (0-50% EtOAc/Hex) to obtain the Boc-protected amine (27 mg, 0.082 mmol, 43%) as a white solid. LC/MS [M+H]+ m/z calc. 331.19, found 331.1. 1H NMR (300 MHz, CDCl_3_) δ 7.78 (d, J = 8.3 Hz, 2H), 7.59 (d, J = 8.7 Hz, 2H), 6.32 (s, 1H), 4.63 (s, 1H), 3.50 (td, J = 7.0, 5.7 Hz, 2H), 3.23 (s, 1H), 3.18 (q, J = 6.5 Hz, 2H), 1.70 (d, J = 7.5 Hz, 2H), 1.62 – 1.52 (m, 2H), 1.46 (s, 11H). This Boc-protected amine (27 mg, 0.082 mmol) was then dissolved in DCM (1 mL) and TFA (0.5 mL) was added. After stirring at rt for 2h, the mixture was diluted in DCM and evaporated repeatedly to remove volatiles and provide the amineTFA salt as an orange oil (32 mg, 0.096 mmol, 117%), which was used without further purification. LC/MS [M+H]^+^ m/z calc. 231.14, found 231.1. 1H NMR (400 MHz, DMSO) δ 8.55 (t, J = 5.7 Hz, 1H), 7.84 (d, J = 8.2 Hz, 2H), 7.63 (s, 2H), 7.57 (d, J = 8.1 Hz, 2H), 4.39 (s, 1H), 3.26 (q, J = 6.6 Hz, 2H), 2.83 – 2.74 (m, 2H), 1.62 – 1.48 (m, 4H), 1.40 – 1.32 (m, 2H).

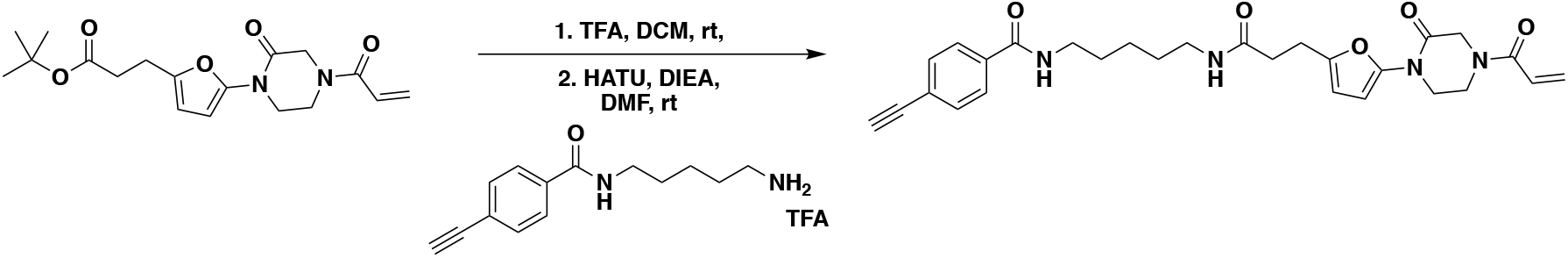

***N*-(5-(3-(5-(4-acryloyl-2-oxopiperazin-1-yl)furan-2-yl)propanamido)pentyl)-4-ethynylbenzamide (NJH-2-075):** tert-butyl 3-(5-(4-acryloyl-2-oxopiperazin-1-yl)furan-2-yl)propanoate (20 mg, 0.057 mmol) was dissolved in DCM (0.5 mL) and treated with TFA (0.25 mL). The mixture was stirred at rt for 45 minutes until the starting material was consumed, followed by dilution with DCM and evaporation to remove volatiles. The carboxylic acid was then dissolved in DMF, and N-(5-aminopentyl)-4-ethynylbenzamide TFA (22 mg, 0.062 mmol), DIEA (50 μL, 0.29 mmol), and HATU (43 mg, 0.11 mmol) were added. After stirring the mixture at rt for 1 h, water was added. The resulting suspension was extracted three times with DCM. Combined organic extracts were washed brine and dried over Na_2_SO_4_, concentrated, and the crude residue was purified by silica gel chromatography (0-4% MeOH/DCM) to obtain NJH-2-075 (7.6 mg, 0.016 mmol, 27%) as a pale yellow oil. HRMS [M+H]^+^ m/z calc. 380.1586, found 380.1581. 1H NMR (300 MHz, CDCl_3_) δ 7.82 (d, J = 8.3 Hz, 2H), 7.58 (d, J = 8.3 Hz, 2H), 6.77 – 6.50 (m, 2H), 6.43 (dd, J = 16.7, 2.1 Hz, 1H), 6.24 (d, J = 3.2 Hz, 1H), 6.06 (d, J = 3.3 Hz, 1H), 5.93 (s, 1H), 5.86 (dd, J = 10.1, 2.1 Hz, 1H), 4.44 (d, J = 17.4 Hz, 2H), 4.01 (s, 2H), 3.91 – 3.84 (m, 2H), 3.46 (q, J = 6.6 Hz, 2H), 3.32 – 3.19 (m, 3H), 2.93 (t, J = 7.2 Hz, 2H), 2.50 (t, J = 7.3 Hz, 2H), 1.72 – 1.61 (m, 2H), 1.60 – 1.46 (m, 2H), 1.44 – 1.35 (m, 2H). 13C NMR (151 MHz, DMSO) δ 171.0, 165.7, 164.6, 150.1, 135.2, 132.1, 128.8, 127.9, 124.7, 106.9, 100.5, 83.4, 83.1, 38.9, 33.8, 29.3, 29.2, 24.3, 24.0.

## Extended Data

**Extended Data Figure 1.**
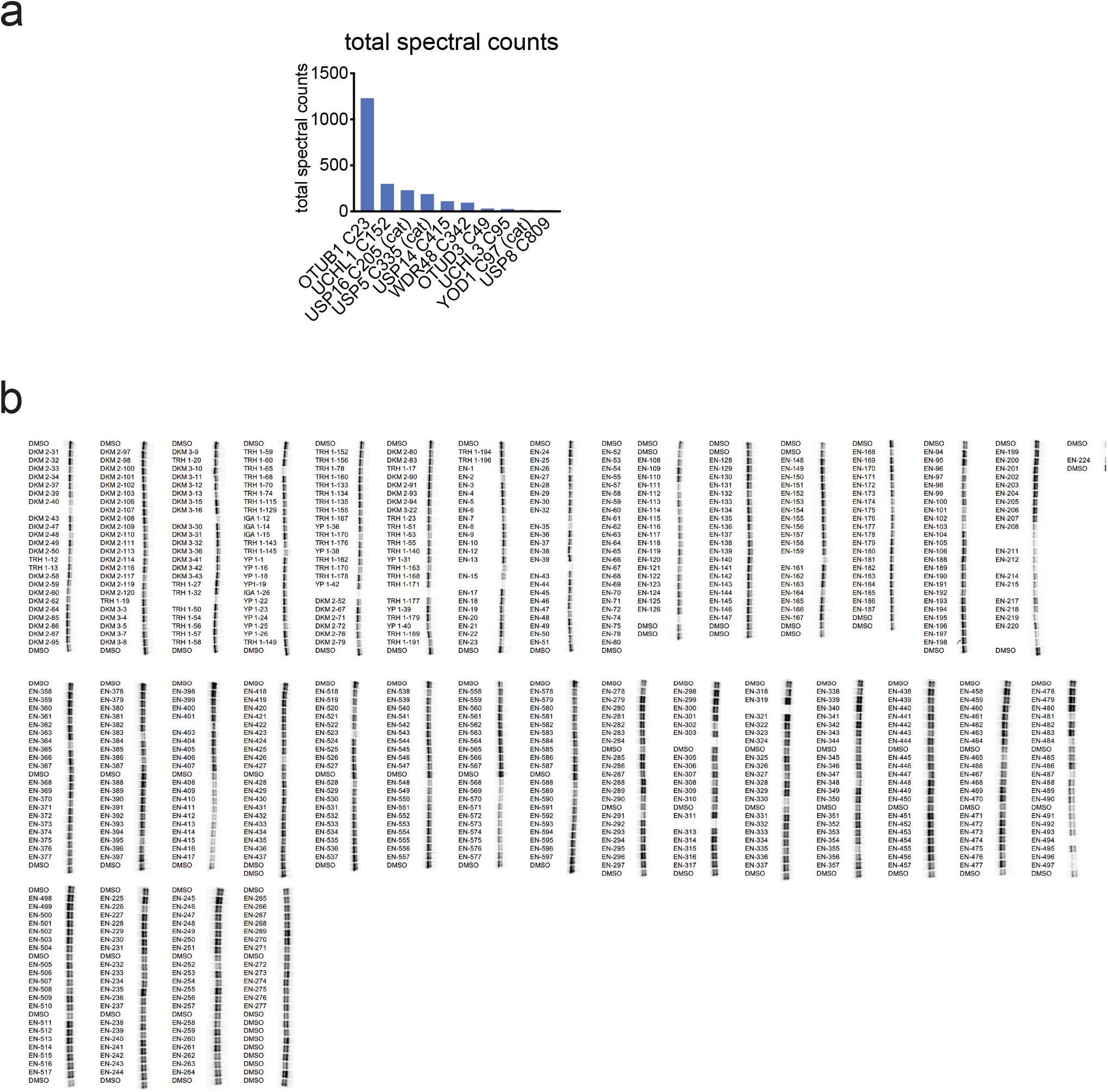
Primary covalent ligand screen against OTUB1. **(a)** Analysis of aggregate chemoproteomic data for DUBs. Top 10 candidate DUBs described in **Figure 1c** for total aggregate spectral counts of the particular probe-modified cysteine found in our aggregate chemoproteomic data showing OTUB1 C23 appears far more frequently in chemoproteomic datasets compared to the other DUBs. **(b)** Covalent ligand screen of cysteine-reactive libraries competed against IA-rhodamine labeling of recombinant OTUB1 to identify binders to OTUB1 by gel-based ABPP. Vehicle DMSO or cysteine-reactive covalent ligands (50 μM) were pre-incubated with OTUB1 for 30 min at room temperature prior to IA-rhodamine labeling (500 nM, 30 min room temperature). OTUB1 was then separated by SDS/PAGE and in-gel fluorescence was assessed and quantified.

**Extended Data Figure 2.**
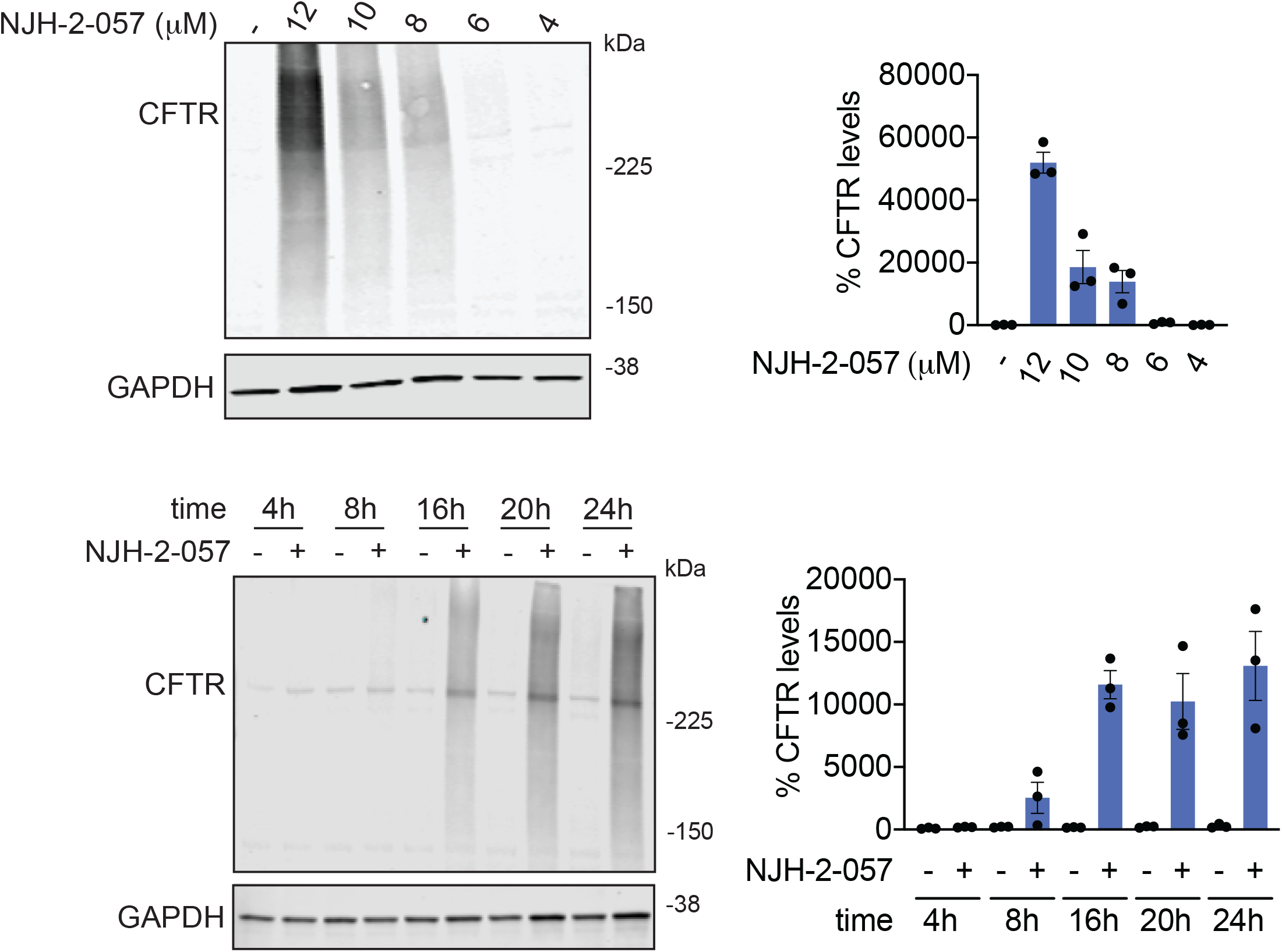
Effect of DUBTACs on mutant CFTR levels. CFBE41o-4.7 cells expressing ΔF508-CFTR were treated with vehicle DMSO or NJH-2-057 and CFTR and loading control GAPDH levels were assessed by Western blotting. For dose-response studies, NJH-2-057 was treated for 24 h. For time-course studies, NJH-2-057 was treated at 10 μM. Dose-response and time-course data gels are representative of n=3 biologically independent samples/group and are quantified in the bar graphs to the right. Data in bar graphs show individual biological replicate values and average ± sem from n=3 biologically independent samples/group.

**Extended Data Figure 3.**
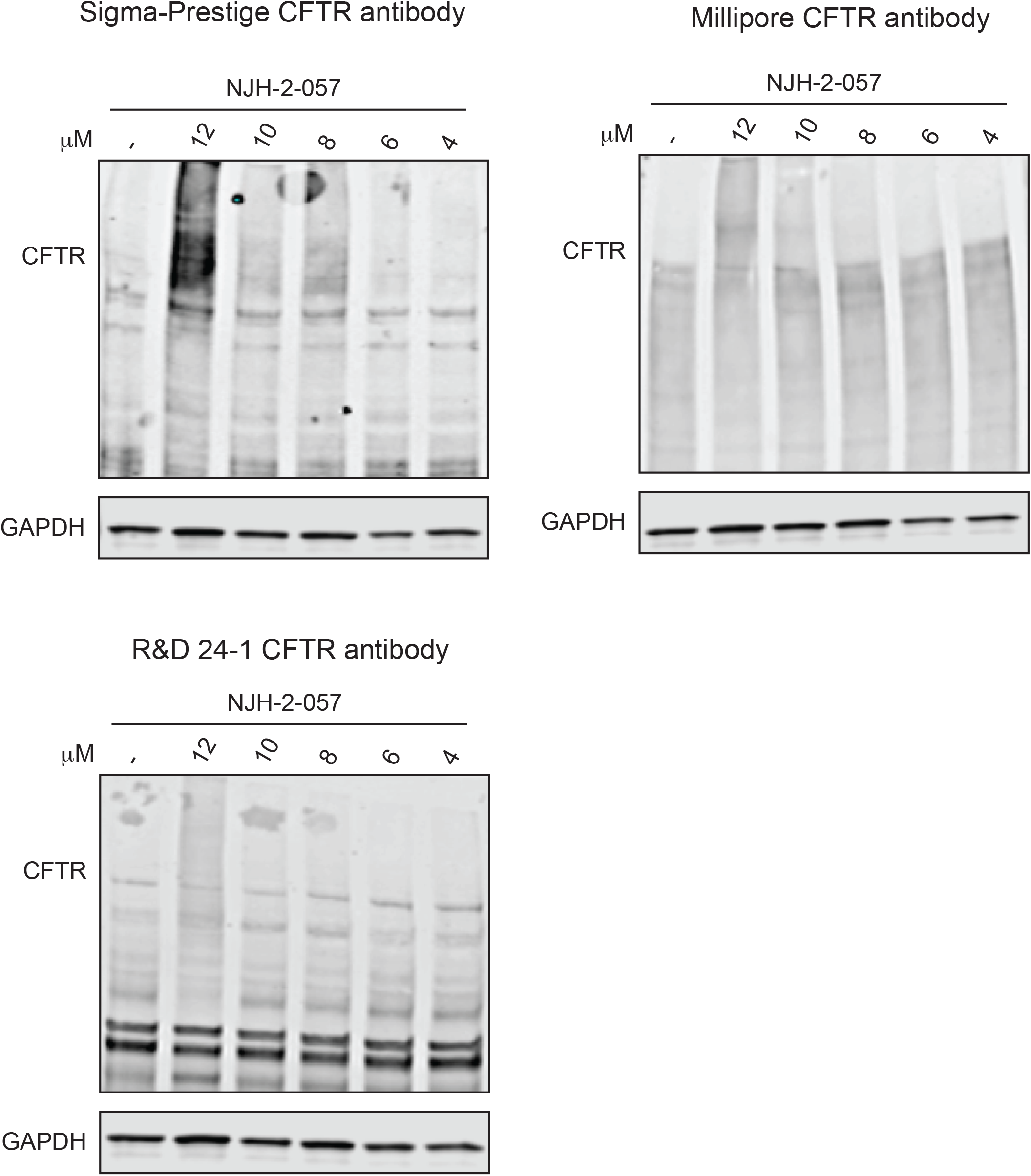
Effect of DUBTACs on mutant CFTR levels. CFBE41o-4.7 cells expressing ΔF508-CFTR were treated with vehicle DMSO or NJH-2-057 and CFTR and loading control GAPDH levels were assessed by Western blotting using three different antibodies against CFTR from the ones used for the main figures. NJH-2-057 was treated for 24 h. Gels are representative of n=3 biologically independent samples/group.

**Extended Data Figure 4.**
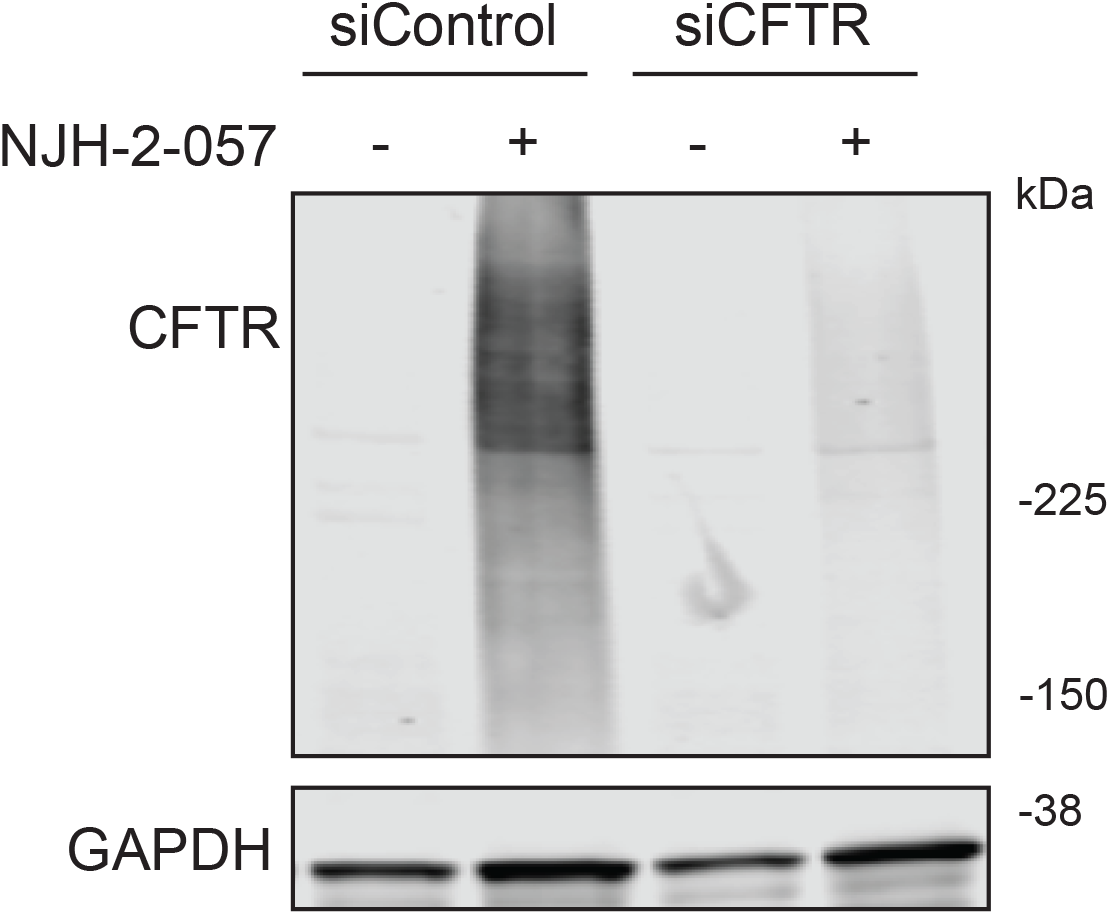
Effect of DUBTACs on mutant CFTR levels in siControl and siCFTR cells. CFBE41o-4.7 cells expressing ΔF508-CFTR were treated with vehicle DMSO or NJH-2-057 (10 μM) for 24 h and CFTR and loading control GAPDH levels were assessed by Western blotting. Blot is representative of n=3 biologically independent samples/group.

**Extended Data Figure 5.**
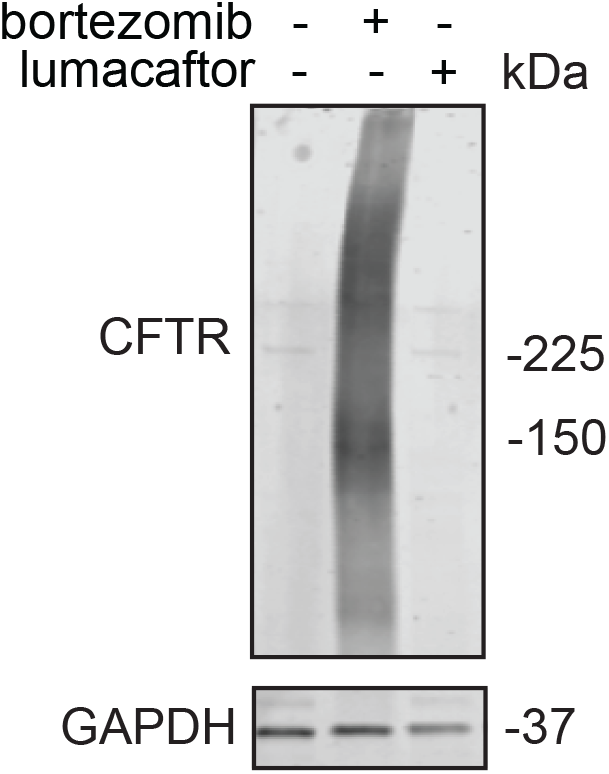
Effect of bortezomib and lumacaftor on mutant CFTR levels. CFBE41o-4.7 cells expressing ΔF508-CFTR were treated with vehicle DMSO, bortezomib (1 μM), or lumacaftor (1 μM) for 24 h and CFTR and loading control GAPDH levels were assessed by Western blotting. The gel shown is representative of n=3 biologically independent samples/group.

**Extended Data Figure 6.**
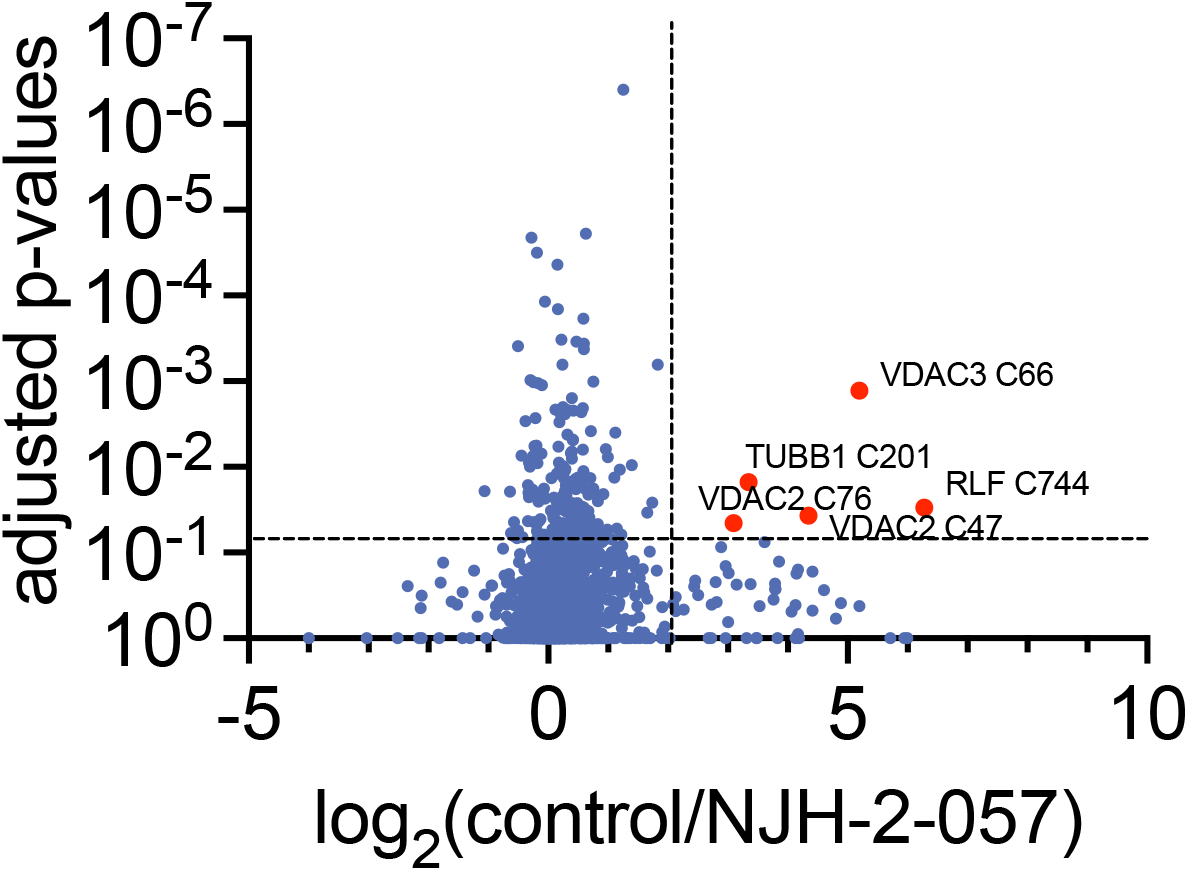
IsoTOP-ABPP analysis of NJH-2-057. CFBE41o-4.7 cells expressing ΔF508-CFTR were treated with vehicle DMSO or NJH-2-057 for 8 h. Resulting cell lysates were labeled with IA-alkyne (200 μM) for 1 h and taken through the isoTOP-ABPP procedure. Shown in red are the probe-modified peptides that showed isotopically light/heavy or control/NJH-2-57 ratios >4 with adjusted p-values <0.05. The data are from n=3 biologically independent samples/group. The full isoTOP-ABPP dataset can be found in **Table S3.**

## References

1. Dixon, S. J. & Stockwell, B. R. Identifying druggable disease-modifying gene products. Curr. Opin. Chem. Biol. 13, 549–555 (2009).

2. Spradlin, J. N., Zhang, E. & Nomura, D. K. Reimagining Druggability Using Chemoproteomic Platforms. Acc. Chem. Res. 54, 1801–1813 (2021).

3. Nalawansha, D. A. & Crews, C. M. PROTACs: An Emerging Therapeutic Modality in Precision Medicine. Cell Chem. Biol. 27, 998–1014 (2020).

4. Burslem, G. M. & Crews, C. M. Proteolysis-Targeting Chimeras as Therapeutics and Tools for Biological Discovery. Cell 181, 102–114 (2020).

5. Schapira, M., Calabrese, M. F., Bullock, A. N. & Crews, C. M. Targeted protein degradation: expanding the toolbox. Nat. Rev. Drug Discov. 18, 949–963 (2019).

6. Banik, S. M. et al. Lysosome-targeting chimaeras for degradation of extracellular proteins. Nature 584, 291–297 (2020).

7. Takahashi, D. et al. AUTACs: Cargo-Specific Degraders Using Selective Autophagy. Mol. Cell 76, 797–810.e10 (2019).

8. Yamazoe, S. et al. Heterobifunctional Molecules Induce Dephosphorylation of Kinases–A Proof of Concept Study. J. Med. Chem. 63, 2807–2813 (2020).

9. Siriwardena, S. U. et al. Phosphorylation-Inducing Chimeric Small Molecules. J. Am. Chem. Soc. 142, 14052–14057 (2020).

10. Sabapathy, K. & Lane, D. P. Therapeutic targeting of p53: all mutants are equal, but some mutants are more equal than others. Nat. Rev. Clin. Oncol. 15, 13–30 (2018).

11. Abbas, T. & Dutta, A. p21 in cancer: intricate networks and multiple activities. Nat. Rev. Cancer 9, 400–414 (2009).

12. Li, B. & Dou, Q. P. Bax degradation by the ubiquitin/proteasome-dependent pathway: Involvement in tumor survival and progression. Proc. Natl. Acad. Sci. U. S. A. 97, 3850–3855 (2000).

13. Ward, C. L., Omura, S. & Kopito, R. R. Degradation of CFTR by the ubiquitin-proteasome pathway. Cell 83, 121–127 (1995).

14. Weerapana, E. et al. Quantitative reactivity profiling predicts functional cysteines in proteomes. Nature 468, 790–795 (2010).

15. Backus, K. M. et al. Proteome-wide covalent ligand discovery in native biological systems. Nature 534, 570–574 (2016).

16. Wiener, R., Zhang, X., Wang, T. & Wolberger, C. The mechanism of OTUB1-mediated inhibition of ubiquitination. Nature 483, 618–622 (2012).

17. Nakada, S. et al. Non-canonical inhibition of DNA damage-dependent ubiquitination by OTUB1. Nature 466, 941–946 (2010).

18. Que, L. T., Morrow, M. E. & Wolberger, C. Comparison of Cross-Regulation by Different OTUB1:E2 Complexes. Biochemistry 59, 921–932 (2020).

19. French, M. E., Koehler, C. F. & Hunter, T. Emerging functions of branched ubiquitin chains. Cell Discov. 7, 1–10 (2021).

20. Wiener, R. et al. E2 ubiquitin-conjugating enzymes regulate the deubiquitinating activity of OTUB1. Nat. Struct. Mol. Biol. 20, 1033–1039 (2013).

21. Riordan, J. R. CFTR function and prospects for therapy. Annu. Rev. Biochem. 77, 701–726 (2008).

22. Kanner, S. A., Shuja, Z., Choudhury, P., Jain, A. & Colecraft, H. M. Targeted deubiquitination rescues distinct trafficking-deficient ion channelopathies. Nat. Methods 17, 1245–1253 (2020).

23. Tomati, V. et al. Genetic Inhibition Of The Ubiquitin Ligase Rnf5 Attenuates Phenotypes Associated To F508del Cystic Fibrosis Mutation. Sci. Rep. 5, 12138 (2015).

24. Sondo, E. et al. Pharmacological Inhibition of the Ubiquitin Ligase RNF5 Rescues F508del-CFTR in Cystic Fibrosis Airway Epithelia. Cell Chem. Biol. 25, 891–905.e8 (2018).

25. Lopes-Pacheco, M. CFTR Modulators: The Changing Face of Cystic Fibrosis in the Era of Precision Medicine. Front. Pharmacol. 10, 1662 (2019).

26. Rath, A., Glibowicka, M., Nadeau, V. G., Chen, G. & Deber, C. M. Detergent binding explains anomalous SDS-PAGE migration of membrane proteins. Proc. Natl. Acad. Sci. U. S. A. 106, 1760–1765 (2009).

27. Spradlin, J. N. et al. Harnessing the anti-cancer natural product nimbolide for targeted protein degradation. Nat. Chem. Biol. 15, 747–755 (2019).

28. Zhang, X., Crowley, V. M., Wucherpfennig, T. G., Dix, M. M. & Cravatt, B. F. Electrophilic PROTACs that degrade nuclear proteins by engaging DCAF16. Nat. Chem. Biol. 15, 737–746 (2019).

29. Zhang, X. et al. DCAF11 Supports Targeted Protein Degradation by Electrophilic Proteolysis-Targeting Chimeras. J. Am. Chem. Soc. 143, 5141–5149 (2021).

30. Ward, C. C. et al. Covalent Ligand Screening Uncovers a RNF4 E3 Ligase Recruiter for Targeted Protein Degradation Applications. ACS Chem. Biol. 14, 2430–2440 (2019).

31. Gavathiotis, E., Reyna, D. E., Bellairs, J. A., Leshchiner, E. S. & Walensky, L. D. Direct and selective small-molecule activation of proapoptotic BAX. Nat. Chem. Biol. 8, 639–645 (2012).

32. Pryde, D. C. et al. The discovery of potent small molecule activators of human STING. Eur. J. Med. Chem. 209, 112869 (2021).

33. Ramanjulu, J. M. et al. Design of amidobenzimidazole STING receptor agonists with systemic activity. Nature 564, 439–443 (2018).

34. Zorn, J. A. & Wells, J. A. Turning enzymes ON with small molecules. Nat. Chem. Biol. 6, 179–188 (2010).

35. Chen, L., Liu, S. & Tao, Y. Regulating tumor suppressor genes: post-translational modifications. Signal Transduct. Target. Ther. 5, 1–25 (2020).

36. Grossman, E. A. et al. Covalent Ligand Discovery against Druggable Hotspots Targeted by Anti-cancer Natural Products. Cell Chem. Biol. 24, 1368–1376.e4 (2017).

37. Bateman, L. A. et al. Chemoproteomics-enabled covalent ligand screen reveals a cysteine hotspot in reticulon 4 that impairs ER morphology and cancer pathogenicity. Chem. Commun. Camb. Engl. 53, 7234–7237 (2017).

38. Roberts, A. M. et al. Chemoproteomic Screening of Covalent Ligands Reveals UBA5 As a Novel Pancreatic Cancer Target. ACS Chem. Biol. 12, 899–904 (2017).

39. Counihan, J. L., Wiggenhorn, A. L., Anderson, K. E. & Nomura, D. K. Chemoproteomics-Enabled Covalent Ligand Screening Reveals ALDH3A1 as a Lung Cancer Therapy Target. ACS Chem. Biol. 13, 1970–1977 (2018).

40. Berdan, C. A. et al. Parthenolide Covalently Targets and Inhibits Focal Adhesion Kinase in Breast Cancer Cells. Cell Chem. Biol. 26, 1027–1035.e22 (2019).

41. Spradlin, J. N. et al. Harnessing the anti-cancer natural product nimbolide for targeted protein degradation. Nat. Chem. Biol. 15, 747–755 (2019).

42. Chung, C. Y.-S. et al. Covalent targeting of the vacuolar H+-ATPase activates autophagy via mTORC1 inhibition. Nat. Chem. Biol. 15, 776–785 (2019).

43. Isobe, Y. et al. Manumycin polyketides act as molecular glues between UBR7 and P53. Nat. Chem. Biol. 16, 1189–1198 (2020).

44. Boike, L. et al. Discovery of a Functional Covalent Ligand Targeting an Intrinsically Disordered Cysteine within MYC. Cell Chem. Biol. 28, 4–13.e17 (2021).

45. Luo, M. et al. Chemoproteomics-enabled discovery of covalent RNF114-based degraders that mimic natural product function. Cell Chem. Biol. (2021) doi:10.1016/j.chembiol.2021.01.005.

46. Xu, T. et al. ProLuCID: An improved SEQUEST-like algorithm with enhanced sensitivity and specificity. J. Proteomics 129, 16–24 (2015).

47. Käll, L., Canterbury, J. D., Weston, J., Noble, W. S. & MacCoss, M. J. Semi-supervised learning for peptide identification from shotgun proteomics datasets. Nat. Methods 4, 923–925 (2007).

